# Selective regulation of transsynaptic alignment and postsynaptic assembly by a novel NCAM family synaptic adhesion molecule

**DOI:** 10.64898/2026.03.13.710942

**Authors:** Paola Van der Linden Costello, Marina N. Wennerberg, Jerrik A. Rydbom, Scott Gratz, Lauren F. Fennema, Kate M. O’Connor-Giles, Heather T. Broihier

## Abstract

Synapse formation underlies the organization of neurons into functional circuits during brain development and requires precise alignment and maturation of pre-and postsynaptic compartments. Many synaptogenic adhesion molecules have been identified that drive target recognition and sustained adhesion between appropriate synaptic partners. Yet the degree to which individual molecules serve selective functions in distinct aspects of synapse formation and maturation remains poorly understood. In particular, how exquisite nanoalignment of pre-and postsynaptic specializations flows from micron-scale adhesive interactions between synaptic partners remains a key unanswered question. Here we shed new light on this question by establishing a specialized set of synaptic functions for *Epithelial limiter of Fasciclin II function (Elff*), an NCAM family member with previously unknown roles in the nervous system. Our structural and functional studies at the glutamatergic Drosophila NMJ indicate that Elff is required for postsynaptic assembly, maturation, and transsynaptic alignment; however, it is not required for presynaptic function, bouton formation, or developmental expansion of the NMJ. Notably, NMJs in *elff* null mutants display reduced glutamate receptor clustering beginning at the embryonic stage when NMJ synapses first form. These poorly defined postsynaptic specializations are frequently out of register with presynaptic release sites, disrupting neurotransmission. Unexpectedly, the striking defects in *elff* nulls occur in the context of both normal active zone number and developmental expansion of the NMJ. These findings suggest a surprising degree of specialization among transsynaptic adhesion complexes and demonstrate that Elff-mediated signaling is critical for the transsynaptic nanoarchitecture of glutamatergic synapses.

## Introduction

A dizzying array of trans-synaptic adhesion molecules shape synapse formation, stabilization, maturation and remodeling. Key molecules, including Neurexins, Neuroligins and Teneurins, are implicated in transsynaptic adhesion, subsynaptic organization, and synaptic differentiation. For example in Drosophila, loss of Neurexin, Neuroligin, or Teneurin results in small neuromuscular junctions (NMJs) with fewer boutons in addition to impaired baseline neurotransmission (Li et al., 2007; Banovic et al., 2010; Chen et al., 2010; Mosca et al., 2012; DePew et al., 2019). The gross morphological defects at these mutant NMJs have complicated efforts to assign subsynaptic functions to individual adhesion molecules. In particular, it is unclear how the essential nanocolumnar organization of pre-and postsynaptic specializations emerges from much larger scale adhesive interactions between pre-and postsynaptic cells (Tang et al., 2016; Biederer et al., 2017).

Neural Cell Adhesion Molecule (NCAM) homologs are conserved synaptic adhesion molecules with widespread roles in axon fasciculation as well as synapse structure and function (Lin et al., 1994a; Rafuse et al., 2000; Ashley et al., 2005; Shetty et al., 2013; Duncan et al., 2021). Fasciclin II (Fas II) is the Drosophila homolog of mammalian NCAMs (Lin et al., 1994a). Both murine NCAM1 and Drosophila Fas II are homophilic adhesion molecules with five immunoglobulin domains (IgSF) and two fibronectin type 3 (FN3) domains. In mice lacking all NCAM1 isoforms, NMJs form but are small and fail to mature (Polo-Parada et al., 2001); (Rafuse et al., 2000). Similarly, NMJs form in *fas II* nulls, but fail to expand, and the animals die at the first-instar larval stage (Schuster et al., 1996a). NCAM family members are also linked to pre-and postsynaptic maturation (Sytnyk et al., 2006; Kohsaka et al., 2007); (Polo-Parada et al., 2001); however, they have not previously been implicated in transsynaptic nanocolumn organization.

Lending clinical significance to the study of NCAM family members at synapses, their dysfunction is well-linked to neurological disorders. Multiple studies implicate NCAM1 or its polysialylated form PSA-NCAM1 to aberrant synaptic plasticity observed in Autism Spectrum Disorder and Schizophrenia (Aonurm-Helm et al., 2016; Williams et al., 2020; Yu et al., 2025). Strikingly, injection of anti-NCAM1 antibodies from individuals with schizophrenia leads to schizophrenia-like synapse and behavior changes in mice, arguing that schizophrenia can be caused by anti-NCAM1 autoantibodies (Shiwaku et al., 2022). Moreover, a recent large-scale multi-omic study identified NCAM1 as a top potential causal gene for PTSD (Nievergelt et al., 2024). Additionally, loss of human NCAM2 is linked to a neurodevelopmental disorder caused by 21q21 deletion (Petit et al., 2015), while its overexpression is associated with Down Syndrome (Parcerisas et al., 2020, 2021).

NCAM orthologs are alternatively spliced, with transmembrane and GPI-linked isoforms (Cunningham et al., 1987; Neuert et al., 2019). Clear evidence exists for cell type-specific roles of distinct isoforms (Hata et al., 2007, 2018; Neuert et al., 2019). Introducing further complexity, additional NCAM family members also regulate neuronal function (Butler et al., 1997; Noordermeer et al., 1998; Babu et al., 2011; Özkan et al., 2013; Fedotov et al., 2018). In *C. elegans*, the IgSF-FN3 domain GPI-linked protein Rig-3 collaborates with the Wnt pathway to antagonize activity-induced changes in neurotransmitter receptor abundance (Babu et al., 2011). In Drosophila, two additional NCAM family members, CG15630/Factor of interpulse interval (Fipi) and CG33543/Epithelial Limiter of Fasciclin II function (Elff) bind Fas II when overexpressed in cultured cells (Özkan et al., 2013). Fipi is required in olfactory sensory neurons for proper male courtship song (Fedotov et al., 2014, 2018). Neuronal roles for Elff have not previously been reported, though its function in wing development was recently described. In the wing, Elff and Fipi both repress EGF receptor activity via both FasII-independent and dependent mechanisms (Garcia-Alonso, 2024). We initially became interested in Elff due to its extracellular domain structure, physical interaction with Fas II, and widespread CNS expression profile leading us to hypothesize roles in synapse structure or function.

Here we generated novel genetic reagents to undertake a comprehensive analysis of Elff function at glutamatergic synapses at the fly NMJ. Surprisingly, we did not detect defects in target recognition, active zone formation or NMJ expansion in *elff* mutants; however, we found clear defects in the assembly and maturation of the postsynaptic compartment. These phenotypes are present from late embryogenesis onward and are accompanied by dramatic misalignment of the diminished postsynaptic glutamate receptor clusters to presynaptic release sites. These structural phenotypes are reinforced by an electrophysiological analysis, which reveals reduced postsynaptic responsiveness in *elff* nulls. We propose that Elff is a critical component of adhesion complexes with essential and specific roles in postsynaptic assembly and maturation as well as transcellular nanoalignment at glutamatergic synapses.

## Results

### The adhesion protein CG33543/Elff localizes to synapses throughout the CNS

The Drosophila genome codes for more than 200 unique adhesion proteins (Özkan et al., 2013), many of which are expressed in the developing and/or adult nervous system. Of the major structural families, IgSF proteins are the most numerous; moreover, their binding partners have been systematically assessed (Özkan et al., 2013), providing key insight into the pathways through which they regulate nervous system structure and function. We noted that two relatively uncharacterized IgSF proteins, CG33543 (Elff) and CG15630 (Fipi), are putative extracellular heterophilic binding partners of the quintessential neuronal adhesion protein Fasciclin II (Fas II). Interestingly, neither Fipi nor Elff are expressed during embryonic stages when Fas II promotes axon outgrowth and guidance (Graveley et al., 2011), arguing for at least some degree of independence among Fas II, Elff, and Fipi. Here, we focus on CG33543/Elff, given its widespread neuronal expression pattern at later developmental stages and in adults (see below).

While Fas II undergoes extensive alternative splicing generating both transmembrane and GPI-linked isoforms, the CG33543 locus is predicted to encode a single GPI-linked isoform containing three IgSF domains and a single Fibronectin 3 (FN3) domain (Figure 1A) (Öztürk-Çolak et al., 2024). CG33543 was recently named *elff* (epithelial limiter of Fasciclin 2 function) on the basis of an RNAi phenotype in the wing (Garcia-Alonso, 2024). To rigorously assess Elff function, we generated two independent null alleles via CRISPR-mediated genome engineering. *elff^JR1^*is a 9.4 KB deletion in which the entire *elff* locus is replaced by an attP site (Figure 1B). *elff^ST1^*, on the other hand, contains an insertion immediately downstream of the signal sequence that includes sfGFP and a visible marker (DsRed) flanked by piggyBac inverted terminal repeat sequences. Prior to DsRed removal, the insertion results in early transcriptional termination and is a predicted null allele (Figure 1B). Both alleles were validated via PCR and sequencing (see Methods) and are homozygous viable and fertile.

**Figure 1.**
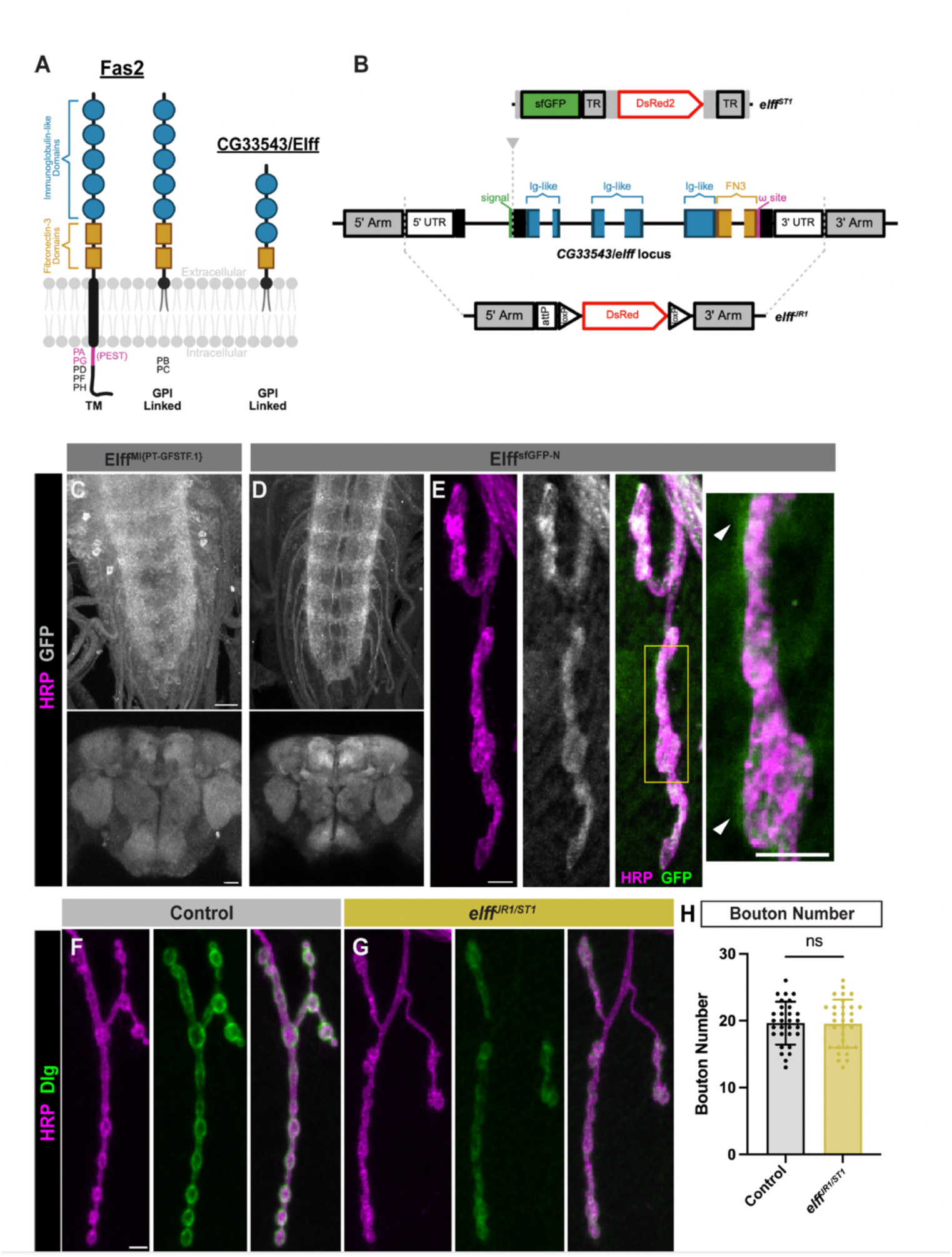
The adhesion protein CG33543/Elff localizes to synapses throughout the CNS. (A) Schematic depicting the predicted protein domain organization of Fas II (left) and CG33543/Elff (right). Fas II has seven known isoforms (PA-PH) containing five IgSF-like domains and two FN3 domains. These Fas II isoforms are predicted to be anchored to the membrane via GPI linkage or a transmembrane (TM) domain with variable cytoplasmic regions. Elff has one known isoform predicted to be GPI-linked with three IgSF domains and one FN3 domain. (B) Schematic view of *elff* allele designs. *elff^JR1^* features a deletion of the entire endogenous coding region, which is replaced with attP and loxP docking sites and a DsRed visible marker. *elff^ST1^* harbors an insertion immediately downstream of the predicted signal sequence, which causes an early stop. (UTR, untranslated region; TR, PiggyBac terminal repeats). (C-D) Representative Z-projections of Drosophila larval ventral nerve cords (top) and adult brains (bottom) from Elff^MI{PT-GFSTF.1}^(C) and Elff^sfGFP-N^ (D) stained for GFP (grayscale). Scale bars: 20 µm. (E) Representative Z-projections of Drosophila muscle 4 NMJs from Elff^sfGFP-N^ larvae stained for GFP (grayscale) and HRP (magenta). Arrowheads indicate Elff^sfGFP-N^ signal outside presynaptic HRP labeling. Scale bar: 5 µm. (F-G) Representative Z-projections of Drosophila NMJs at muscle 4 from the indicated genotypes stained for Dlg (green) and HRP (magenta). Scale bar: 5 µm. (H) Quantification of bouton number. Data are mean ± SD (control: 19.6 ± 3.2, *elff^JR1/ST1^*: 19.6 ± 3.6). Significance determined by unpaired t-test [ns, not significant]. n ≥ 30, Animals ≥ 8.

Following PiggyBac transposase-mediated removal of the visible marker, *elff^ST1^* generates an endogenous N-term epitope tagged protein (Elff^sfGFP-N^). We characterized Elff expression in the nervous system using both Elff^sfGFP-N^ and a MIMIC protein trap, Elff^MI{PT-GFSTF.1}^ (Nagarkar-Jaiswal et al., 2015). Both Elff^MI{PT-GFSTF.1}^ and Elff^sfGFP-N^ are strongly expressed in synaptic neuropil in third-instar larvae and the adult brain (Figure 1C-D). Additionally, Elff^sfGFP-N^ is present at the third-instar larval NMJ, where it strongly overlaps with the neuronal membrane marker HRP and is also observed surrounding presynaptic boutons (Figure 1E), suggesting Elff resides in both the pre-and post-synaptic compartment at the NMJ. We conclude that Elff is broadly synapse-associated during development and in adults.

As a first test of a functional requirement, we investigated if Elff is required for NMJ growth. We characterized the phenotype of animals transheterozygous for the two independent alleles (*Elff^JR1/ST1^*) to minimize potential off-target effects of CRISPR-mediated mutagenesis. We find no difference in the number of synaptic boutons in *Elff^JR1/ST1^* mutant NMJs relative to controls (Figure 1F-H), arguing that Elff does not control baseline NMJ growth. Both the viability and the lack of a gross morphological phenotype in these mutants are in stark contrast to the early larval lethality and NMJ growth defects in Fas II mutations (Lin et al., 1994b; Schuster et al., 1996a, 1996b; Davis et al., 1997; Yoshihara et al., 1997), arguing that Elff does not partner with Fas II to control motor axon targeting or overall NMJ adhesion. Moreover, loss of Elff does not alter overall Fas II levels at the NMJ (Figure S1), indicating that Elff is not essential for expression of Fas II at the NMJ.

### Elff is necessary for postsynaptic maturation at the NMJ

The Drosophila equivalent of the postsynaptic density (PSD) at the NMJ is the subsynaptic reticulum (SSR), which is a weblike structure of filamentous components (Atwood et al., 1993). In the preceding analysis, we co-labeled NMJs with HRP and the SSR marker Dlg (Drosophila PSD95) to distinguish Type 1b and 1s boutons (Zito et al., 1999; Menon et al., 2013) and noticed that *elff* mutant NMJs display an apparent loss of postsynaptic Dlg (Figure 1F-G). Indeed, quantification revealed an approximate 50% decrease in average Dlg intensity in *elff^JR1/ST1^* homozygotes (Figure 2A-B, E). We found a similar decrease in Dlg intensity in homozygous *elff^sfGFP-N^* animals (Figure 2E), indicating that the N-terminal GFP tag unfortunately interferes with Elff function. In contrast, Dlg expression levels are normal in animals homozygous for the MIMIC protein trap (Figure 2E), indicating that Elff function is preserved in this background. To probe specific cellular requirements of *elff*, we employed two independent RNAi lines for cell-type specific knockdown. Motoneuron-or muscle-specific *elff* knockdown via at least one RNAi line results in a loss of Dlg (Figure 2A, C-E), arguing that Elff is required in both the pre-and postsynaptic cells for proper Dlg expression.

**Figure 2.**
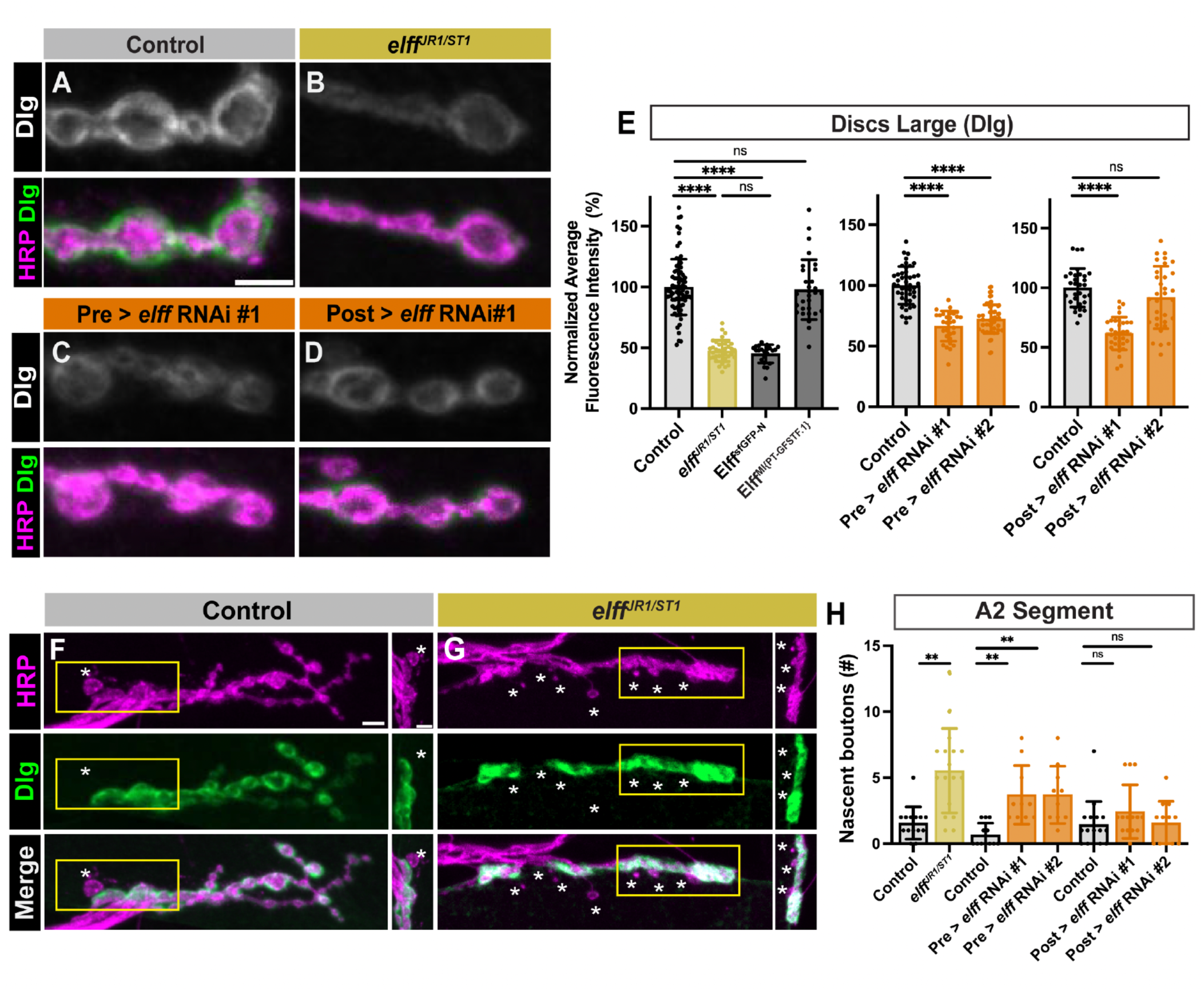
Elff is necessary for post-synaptic maturation at the NMJ. (A-D) Representative Z-projections of NMJ boutons from the indicated genotypes stained for Dlg (green) and HRP (magenta). For clarity, single channels are shown in grayscale next to or below merged images across all figures. For all figures, drivers used for presynaptic (Pre) and postsynaptic (Post) knockdown are D42Gal4 and 24BGal4, respectively. Scale bar: 5 µm. (E) Quantification of Dlg fluorescence intensity. Data are mean values normalized to control ± SD (control: 100.0 ± 22.9, *elff*^JR1/ST1^: 47.5 ± 8.7, Elff^sfGFP-N^: 45.1 ± 7.5, Elff^MI{PT-GFSTF.1}^: 97.8 ± 24.6; control: 100.0 ± 15.5, Pre > *elff* RNAi #1: 65.6 ± 12.4, Pre > *elff* RNAi #2: 72.4 ± 11.8; control: 100.0 ± 16.1, Post > *elff* RNAi #1: 61.8 ± 13.7, Post > *elff* RNAi #2: 92.0 ± 26.1). (F-G) Representative Z-projections of boutons at NMJ 6/7 from the indicated genotypes stained for Dlg (green) and HRP (magenta). Asterisks indicate nascent boutons. Scale bar: 5 µm; inset scale bar: 2 µm. (H) Quantification of nascent boutons. Data are mean ± SD (control: 1.6 ± 1.2, *elff^JR1/ST1^*: 5.5 ± 3.2; control: 0.7 ± 0.9, Pre > *elff* RNAi #1: 3.7 ± 2.2, Pre > *elff* RNAi #2: 3.7 ± 2.2; control: 1.4 ± 1.8, Post > *elff* RNAi #1: 2.4 ± 2.0, Post > *elff* RNAi #2: 1.6 ± 1.6). Significance determined by Kruskal-Wallis test with Dunn’s multiple comparisons test [ns, not significant, **p<0.01, ****p<0.0001]. n ≥ 10, Animals ≥ 8.

If Elff regulates postsynaptic maturation, we hypothesized we would also see an increase in nascent boutons in *elff* homozygotes. Nascent (or ghost) boutons are newly budded boutons lacking a corresponding postsynaptic compartment. They are typically identified as an individual HRP-positive presynaptic bouton in the absence of a corresponding Dlg-positive postsynaptic specialization. They occur with low frequency in wild-type strains and with higher frequency in mutants with impaired postsynaptic maturation (Ataman et al., 2008; Mosca and Schwarz, 2010; McLaughlin et al., 2016). Because Dlg levels are reduced in *elff* mutants, we increased the confocal gain in the Dlg channel of *elff* LOF mutants to determine whether any Dlg is present in apposition to newly formed boutons. In line with our hypothesis, loss of *elff* results in a roughly four-fold increase in nascent boutons (Figure 2F-H). We employed an RNAi-mediated approach to determine cell-type specific requirements for Elff.

Presynaptic *elff* knockdown with either RNAi line results in significantly increased numbers of nascent boutons (Figure 2H). On the other hand, postsynaptic *elff* knockdown results in a trending, but non-significant, increase with one of the RNAi lines (Figure 2H). Taken together, these findings argue that Dlg is particularly sensitive to presynaptic levels of Elff, while postsynaptic Elff may also play a supporting role in promoting proper Dlg expression.

### Elff acts transsynaptically to promote alpha-Spectrin accumulation

The loss of Dlg in *elff* mutants suggests a failure of post-synaptic maturation. To investigate the extent of the defect, we assessed alpha-Spectrin. The Spectrin cytoskeleton is composed of an ordered array of alpha and beta Spectrin heterodimers and is an evolutionarily conserved component of pre-and postsynaptic specializations (Malchiodi-Albedi et al., 1993). At the Drosophila NMJ, the postsynaptic Spectrin scaffold is particularly apparent and is observed as a halo surrounding HRP-positive presynaptic boutons (Figure 3A) Strikingly, we find a greater than 75% reduction in alpha-Spectrin levels in *elff^JR1/ST1^* homozygotes (Figure 3A-B, E), with little postsynaptic Spectrin remaining in *elff* null third-instar larvae. In line with an exclusive presynaptic requirement for Elff, motoneuron-specific knockdown, but not muscle-specific knockdown, leads to reduced alpha-Spectrin (Figure 3A, C-E).

**Figure 3.**
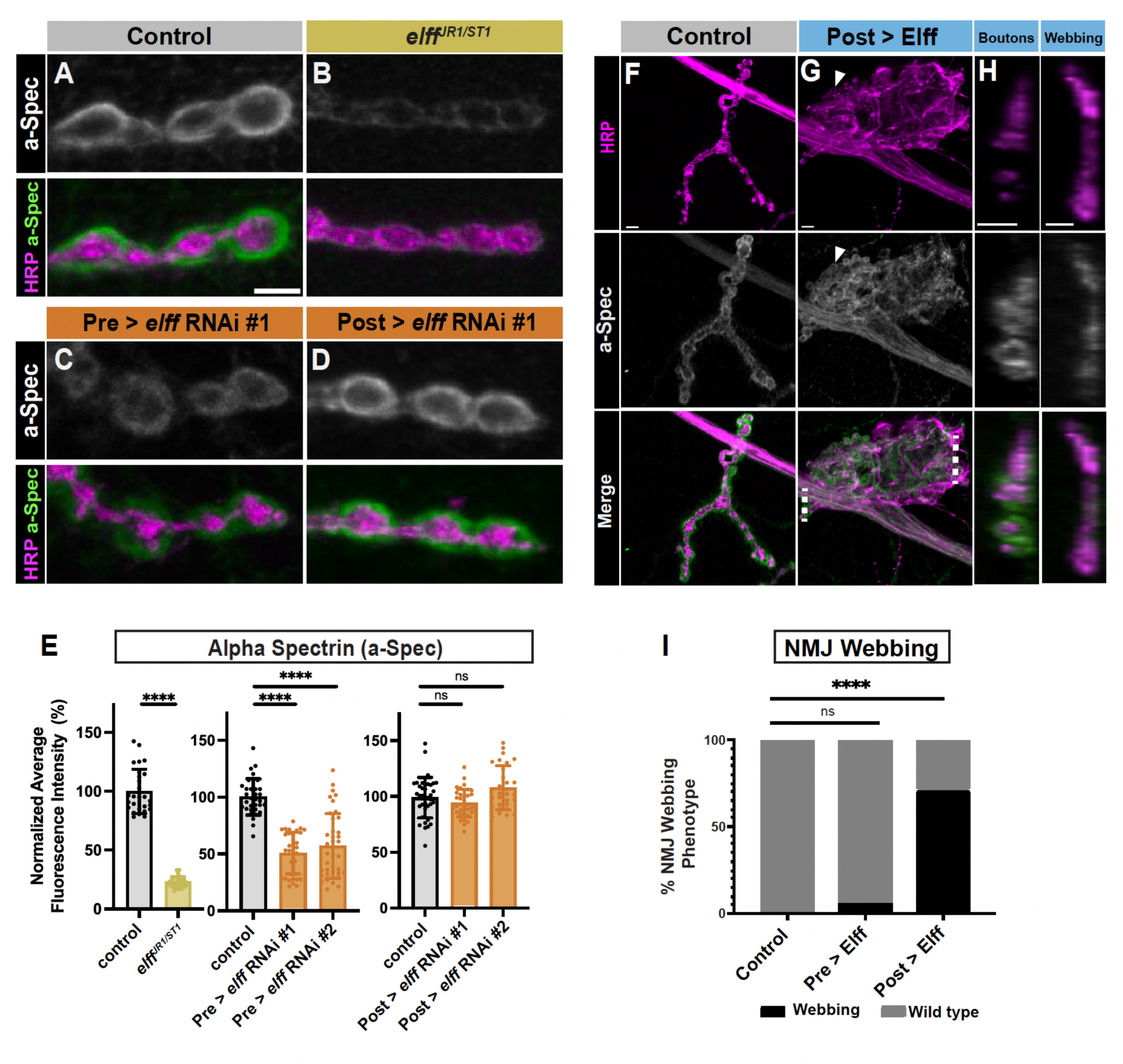
Elff acts transsynaptically to promote alpha-Spectrin accumulation. (A-D) Representative Z-projections of boutons from the indicated genotypes stained for alpha-Spectrin (green) and HRP (magenta). Scale bar: 5 µm. (E) Quantification of alpha-Spectrin intensity. Data are mean values normalized to control ± SD (control: 100.0 ± 18.8, *elff^JR1/ST1^*: 23.1 ± 4.5; control: 100.0 ± 16.0, Pre > *elff* RNAi #1: 50.8 ± 18.4, Pre > *elff* RNAi #2: 57.0 ± 28.5; control: 100.0 ± 17.0, Post > *elff* RNAi #1: 94.1 ± 12.4, Post > *elff* RNAi #2: 107.8 ± 19.8). (F-G) Representative Z-projections of muscle 4 NMJs of the indicated genotypes stained for alpha-Spectrin (green) and HRP (magenta). Arrows indicate synaptic boutons. Scale bars: 5 µm. (H) Orthogonal projections from synaptic boutons (left) and expanded HRP webbing (right) in Post > Elff. Dotted lines in G indicate the areas used to generate orthogonal projections. Scale bars: 5 µm. (I) Quantification of NMJ webbing phenotype (control: 0% of NMJs with webbing, Pre > Elff: 5.50% of NMJs with webbing, Post > Elff: 71.0% of NMJs with webbing). Significance determined by Kruskal-Wallis test with Dunn’s multiple comparisons (E) or chi-square test (I) [ns, not significant, ****p<0.0001]. n ≥ 27, Animals ≥ 8.

These findings indicate that presynaptic Elff promotes alpha-Spectrin expression. In the course of our genetic analysis, we generated UAS-Elff transgenic animals to enable an overexpression analysis.

Unexpectedly, postsynaptic Elff overexpression via 24BGal4 has a widespread effect on NMJ morphology. At roughly 75% of NMJs, postsynaptic Elff overexpression disrupts normal terminal morphology and drives large, lamellipodial-shaped HRP-positive axon terminals, here referred to NMJ webs (Figure 3F-I). We quantified this phenotype at NMJ4, but it is not NMJ-specific and occurs at NMJs on all body wall muscles (data not shown). The normal pattern of motor axon guidance is not altered; the webbing appears over the surface of individual muscles where individual NMJs normally form. Indeed, within the large web-like axon terminals, strings of round boutons are present (arrowheads in Figure 3G). This phenotype is not specific to our UAS-Elff transgene because it is also observed at high frequency with an independently generated UAS-Elff construct (Garcia-Alonso, 2024) (data not shown). Notably, presynaptic Elff overexpression does not have comparable effects (Figure 3I).

The frequent conversions of terminal strings of boutons to enlarged lamellipodial-shaped webs suggests that high levels of postsynaptic Elff are sufficient to drive pervasive cytoskeletal rearrangements on the presynaptic side. Given that presynaptic loss of Elff results in reduced postsynaptic alpha-Spectrin, we wondered whether postsynaptic Elff overexpression results in increased presynaptic alpha-Spectrin. In line with our hypothesis, we find that the enlarged presynaptic HRP-positive webbing contains robust alpha-Spectrin staining (Figure 3G). We conclude that the alpha-Spectrin signal is presynaptic both because it is contiguous with axonal alpha-Spectrin and because it is contained within the HRP signal (Figure 3G). Whereas alpha-Spectrin is predominantly postsynaptic around the small strings of boutons found within the large web, alpha-Spectrin appears entirely within the web itself (Figure 3H). These results indicate that postsynaptic Elff overexpression promotes a large-scale morphological transformation of NMJ terminal arbors from strings of round boutons to enlarged webbed axonal terminals rich with alpha-Spectrin. Together, these complementary loss-and gain-of-function analyses indicate that Elff acts transsynaptically to promote alpha-Spectrin accumulation at the NMJ.

### Elff is necessary for baseline synaptic transmission and larval locomotion

We next tested if impaired postsynaptic maturation observed in *elff* loss-of-function mutants are accompanied by deficits in synaptic function by measuring spontaneous (mEJPs) and evoked (EJPs) junctional potentials from muscle 6 in third-instar *elff* null mutants. mEJP amplitude, a measure of the response of the postsynaptic muscle to fusion of a single glutamatergic vesicle, is reduced by 35% in *elff* null mutants, while mEJP frequency is reduced by more than 60% (Figure 4A-C). The impairment in mEJP frequency could result from either a decrease in presynaptic release probability or decreased postsynaptic responsiveness to the fusion of single neurotransmitter vesicles. To begin to discriminate between these two possibilities, we assessed quantal content, or the number of vesicles released per action potential, which is estimated as the ratio of EJP amplitude to mEJP amplitude. We find that EJP and mEJP amplitudes are similarly reduced in *elff* LOF mutants (Figure 4B, D), leading to equivalent measures of quantal content in control and *elff* mutants (Figure 4E). These data indicate that similar numbers of synaptic vesicles are released in response to a stimulus, implying a largely postsynaptic origin of the physiological defect.

**Figure 4.**
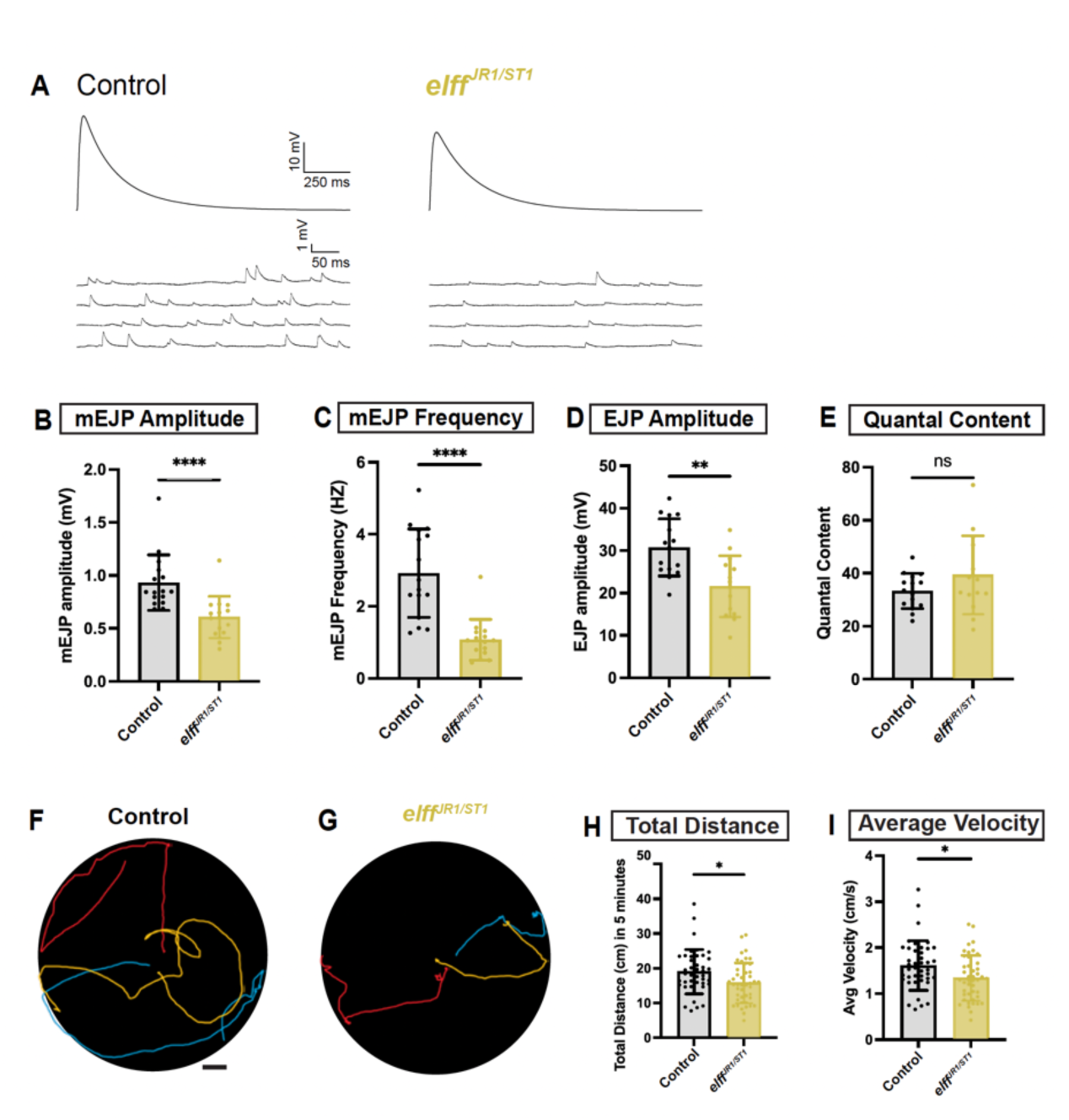
Elff is necessary for baseline synaptic transmission and larval locomotion. (A) Representative traces of evoked (top) and spontaneous (bottom) excitatory junction potentials (EJPs) from the indicated genotypes. Recordings obtained from larval muscle 6 of abdominal segments A3 and A4. (B-E) Quantification of (B) mean mEJP amplitude (control: 0.9 mV ± 0.3, *elff^JR1/ST1^*: 0.6 mV ± 0.2), (C) mean mEJP frequency (control: 2.9 HZ ± 1.2, *elff^JR1/ST1^*: 1.1 HZ ± 0.6), (D) mean EJP amplitude (control: 30.7 mV ± 6.7, *elff^JR1/ST1^*: 21.6 mV ± 7.2), and (E) mean quantal content (control: 33.3 ± 6.6, *elff^JR1/ST1^*: 39.4 ± 14.7). Data are mean ± SD. Each data point represents one animal (n ≥ 14). (F-G) Representative traces of third-instar larval crawling behavior following a 5 min tracking period for animals of the indicated genotypes. Scale bar: 1 cm. (H-I) Quantification of total distance crawled (H) and average velocity (I) obtained from manual tracking (H, control: 19.0 cm ± 6.4, *elff^JR1/ST1^*: 15.8 cm ± 5.8); (I, control: 1.6 cm/s ± 0.5, *elff^JR1/ST1^*: 1.3 cm/s ± 0.5). Each data point represents one animal (n ≥ 42). Significance determined by Mann-Whitney test (B-C) or unpaired t-test (D-E, H-I) [ns, not significant, *p<0.05, **p<0.01, ****p<0.0001].

To ascertain whether the defect in neurotransmission is associated with impaired motor function, we tested whether loss of *elff* results in altered locomotion of third instar larvae. We find that Elff is required for normal motility since its loss leads to significant reductions in both the mean velocity and the total distance travelled by larvae over a five-minute period (Figure 4F-I). Together, functional and behavioral analyses indicate that Elff is specifically required for neurotransmission at the NMJ, and more generally, for robust organismal locomotion.

### Elff is required for glutamate receptor localization at the NMJ

The diminished mEJP amplitude and frequency in *elff* mutants is reminiscent of phenotypes observed in Drosophila ionotropic glutamate receptor (GluR) subunit mutants (Petersen et al., 1997; DiAntonio et al., 1999; Marrus et al., 2004; Featherstone et al., 2005), suggesting that Elff promotes proper GluR levels or localization. Thus, we measured synaptic levels of individual GluR subunits in *elff* mutants using a collection of subunit-specific antibodies (Perry et al., 2017, 2022; Goel and Dickman, 2018; Qiu et al., 2025). At the Drosophila NMJ, glutamate receptors are heterotetrameric complexes comprised of one of two alternate subunits: GluRIIA or GlurIIB, as well as one subunit each of the three essential subunits: GluRIIC, GluRIID, and GluRIIE (Marrus et al., 2004; Qin et al., 2005).

We saw a significant loss of both GluRIIA and GluRIIB from *elff* null NMJs, with a greater than 50% reduction in GluRIIA levels (Figure 5A-F). The loss of both alternative GluR subunits raised the possibility that overall GluR levels are reduced at *elff* mutant NMJs. To investigate this idea, we quantified synaptic levels of GluRIIC and GluRIIE, two of the essential subunits, at control and *elff* mutant NMJs. In line with our hypothesis, intensities of GluRIIC and GluRIIE are also reduced by greater than 50% in *elff* mutants (Figure 5G-L), with individual receptor clusters appearing smaller and difficult to resolve. The decrease in both alternative and essential GluR subunits indicates that overall levels of heterotetrameric complexes are significantly reduced in *elff* null mutants.

**Figure 5.**
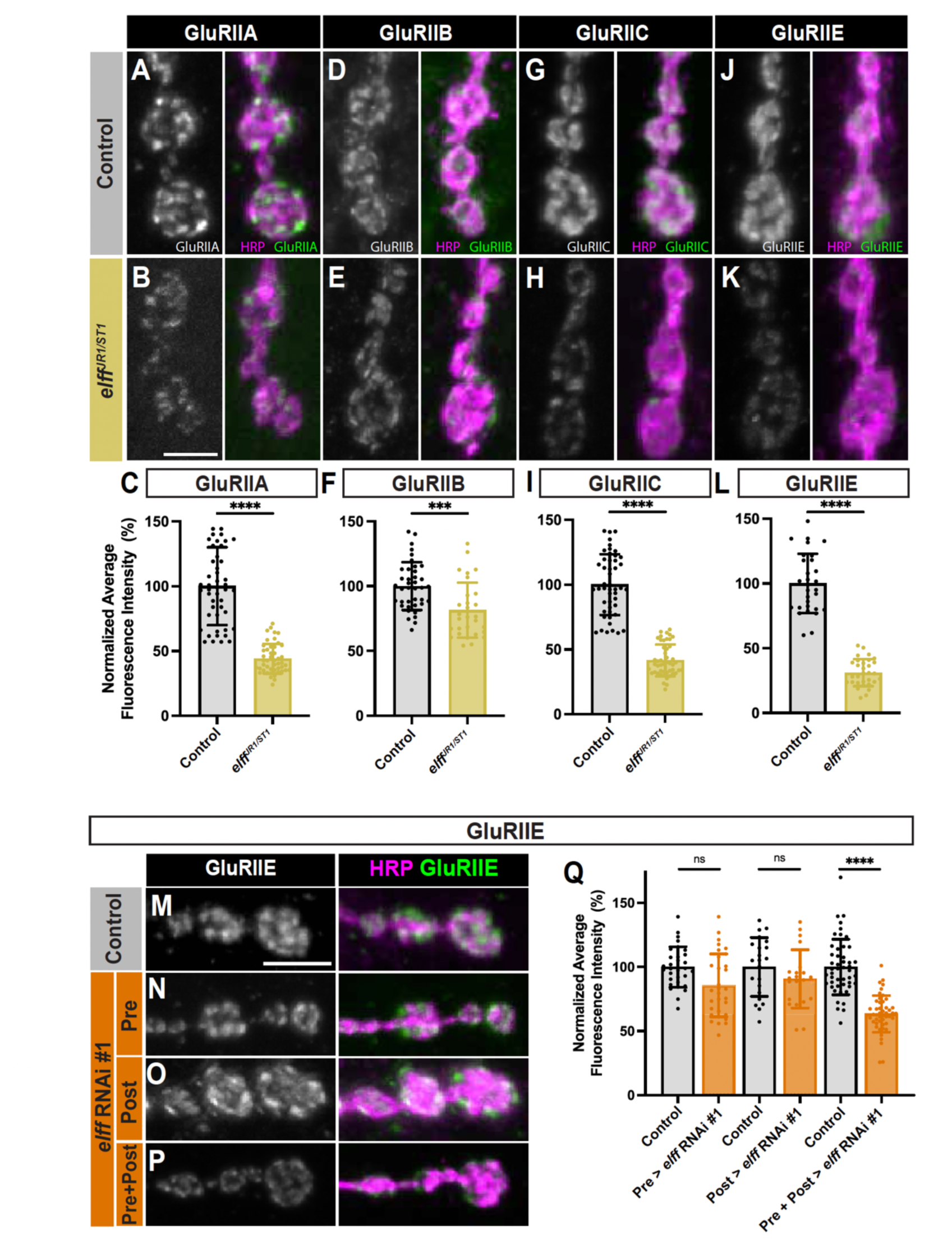
Elff is required for glutamate receptor localization at the NMJ. (A-B, D-E, G-H, J-K) Representative Z-projections of boutons from the indicated genotypes stained for (A-B) GluRIIA, (D-E) GluRIIB, (G-H) GluRIIC, or (J-K) GluRIIE (green) and HRP (magenta). Scale bar: 5 µm. (C, F, I, L) Quantification of fluorescence intensity for (C) GluRIIA (control: 100.0 ± 30.1, *elff^JR1/ST1^*: 44.1 ± 11.4), (F) GluRIIB (control: 100.0 ± 18.5, *elff^JR1/ST1^*: 81.4 ± 21.2), (I) GluRIIC (control: 100.0 ± 23.4, *elff^JR1/ST1^*: 41.6 ± 12.3), and (L) GluRIIE (control: 100.0 ± 23.0, *elff^JR1/ST1^*: 31.1 ± 10.4). Data are mean values normalized to control ± SD. (M-P) Representative Z-projections of boutons from the indicated genotypes. For all figures, drivers for knockdown on both sides of the synapse (Pre + Post) are elavGal4 and 24BGal4. Scale bar: 5 µm. (Q) Quantification of GluRIIE intensity (control: 100.0 ± 15.8, Pre > *elff* RNAi #1: 85.5 ± 24.6; control: 100.0 ± 22.9, Post > *elff* RNAi #1: 90.7 ± 22.8; control: 100.0 ± 21.6, Pre+Post > *elff* RNAi #1: 63.5 ± 14.3). Significance determined by Mann-Whitney test (C, F, I), unpaired t-test (L) or one-way ANOVA with Tukey’s multiple comparisons test (Q) [ns, not significant, ***p<0.001, ****p<0.0001]. n ≥ 21, Animals ≥ 6.

To test whether Elff is required pre-or postsynaptically, we assessed levels of the essential GluR subunit GluRIIE with cell type-specific RNAi-mediated knockdown. We chose to focus on GluRIIE since it is an essential subunit and is particularly clearly reduced in *elff* nulls. We found modest, non-significant, decreases in GluRIIE with either motoneuron-specific or muscle-specific knockdown (Figure 5M-O, Q). These data raise the possibility that Elff expression in either the pre-or postsynaptic cell is sufficient to promote GluR complex localization or stabilization. To test this idea, we simultaneously knocked down *elff* in neurons and muscle cells and found a near 50% reduction in GluRIIE levels at NMJ synapses (Figure 5M, P-Q). These data imply that Elff is present on both sides of the synapse and are in line with the model that Elff can support proper GluR levels if secreted from either the pre-or postsynaptic side. Together, these findings argue that the defects in mini frequency and amplitude in *elff* nulls are the result of reduced postsynaptic GluR clusters and indicate that Elff is broadly required for postsynaptic maturation.

We next probed a possible glial requirement for Elff. As a predicted GPI-linked FN3-IgSF protein, Elff is closely related to Fas II and NCAM GPI-linked isoforms. In both Drosophila and mammalian models, the secreted GPI-linked isoforms are expressed predominantly by glial cells (Maness and Schachner, 2007; Neuert et al., 2019). Indeed, expression of a GPI-linked isoform of Fas II in glia rescues *fas* II null mutants to viability in Drosophila (Neuert et al., 2019). To test whether Elff likewise functions in glia, we tested whether pan-glial *elff* knockdown results in reduced Dlg or GluRIIE expression– two sensitive read-outs of *elff* function. We find that expression of *elff* RNAi via RepoGAL4 fails to alter either Dlg levels or GluRIIE clustering (Figure S2 A-B). Thus, Elff is not required by glia to drive postsynaptic maturation.

### Elff promotes initial assembly of the postsynaptic compartment

The preceding analysis demonstrates that Elff is necessary for multiple facets of postsynaptic maturation at the wandering third instar larval stage, corresponding to roughly five days after egg laying (AEL).

The analysis could not determine whether Elff is required for the initial formation of GluR clusters or rather for their stabilization since NMJ synapses are assembled at the end of embryogenesis, at 24 hours AEL. Thus, an assessment of Elff function in synapse formation demands a phenotypic analysis at earlier developmental stages. Initial synapse formation at stage 17 embryonic NMJs is not often studied in part because standard embryo staining protocols do not effectively stain very late-stage embryos due to the presence of the embryonic cuticle.

We started with the more straightforward task of assessing postsynaptic markers at 48 h AEL, or mid second-instar stage. Strikingly, loss of Elff results in an almost complete loss of alpha-Spectrin and Dlg in L2 stage NMJs (Figure 6A-D, G-H) as well as a near complete absence of GluR complexes marked by anti-GluRIIE (Figure 6E-F, I) in young larvae. The finding that loss of Elff results in more pronounced phenotypes at L2 vs L3 hints that Elff may be required for synapse formation rather than later steps of synapse stabilization or maturation. These data are also consistent with a large body of literature arguing that robust homeostatic mechanisms are in place that can compensate for impaired NMJ function (Dickman and Davis, 2009; Davis, 2013; Perry et al., 2017; Goel and Dickman, 2018; Goel et al., 2019).

**Figure 6.**
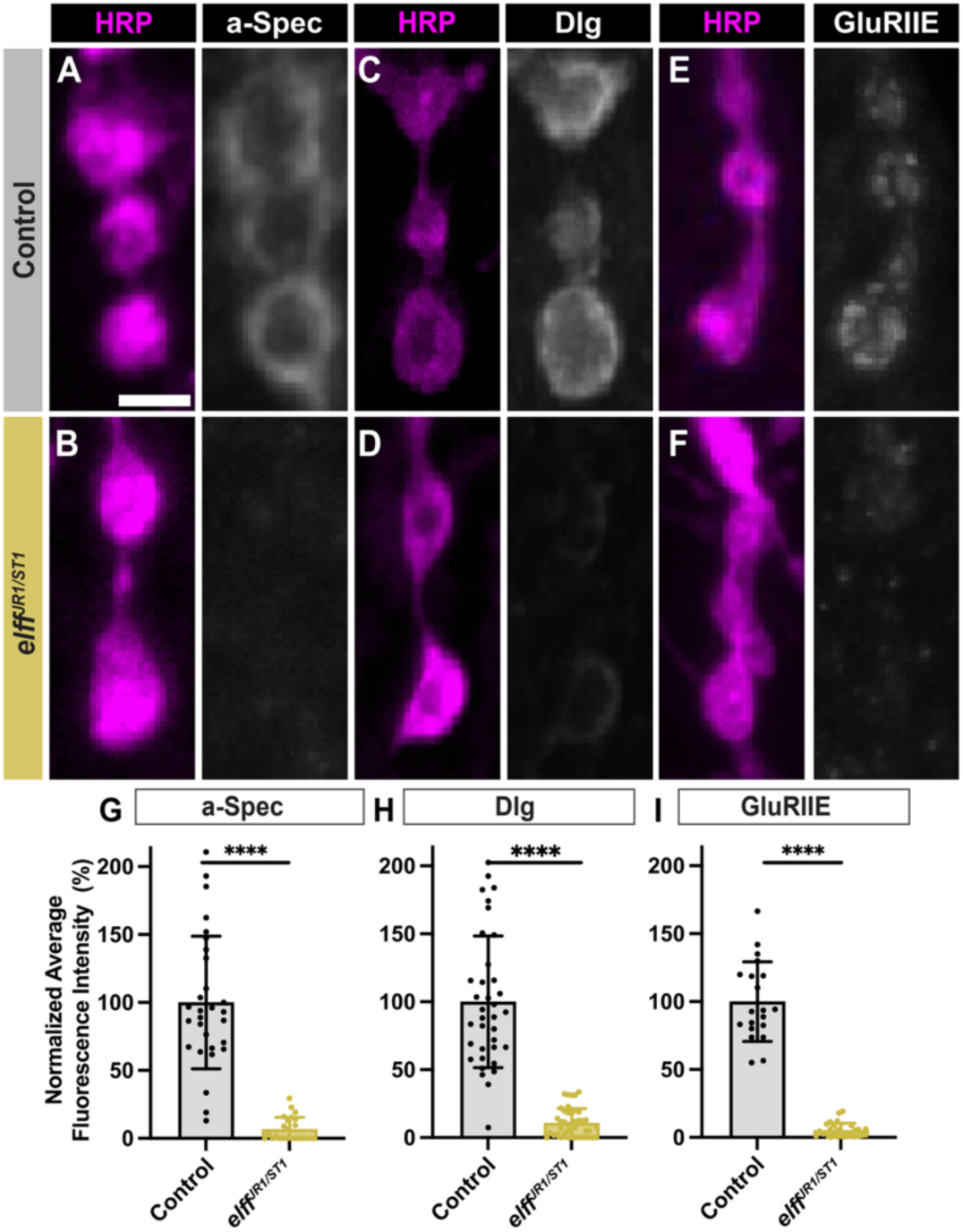
Elff is required for the differentiation of the postsynaptic compartment during mid-larval development. (A-F) Representative Z-projections of boutons in second instar larvae of the indicated genotypes, stained for HRP (magenta) and (A-B) alpha-Spectrin, (C-D) Dlg, or (E-F) GluRIIE (grayscale). Scale bar: 2 µm. (G) Quantification of alpha-Spectrin intensity. Data are mean values normalized to control ± SD (control: 100 ± 48.80, *elff^JR1/ST1^*: 6.665 ± 8.820). (H) Quantification of Dlg intensity (control: 100 ± 48.42, *elff^JR1/ST1^*: 10.81 ± 10.57). (I) Quantification of GluRIIE intensity (control: 100 ± 29.30, *elff^JR1/ST1^*: 5.311 ± 5.195). Significance determined by Mann-Whitney test [****p<0.0001]. n ≥ 20, Animals ≥ 5.

These strong second instar phenotypes prompted us to interrogate initial embryonic synapse formation. To this end, we developed a new stage 17 embryonic staining protocol to better preserve embryonic NMJ morphology and enable a quantitative analysis of active zone and postsynaptic receptor levels and apposition (see Materials and Methods). We analyzed mid-stage 17 embryonic NMJs since this is after motor axon terminals elaborate on their target muscles, form boutons, and initiate synapse formation. At this stage, there are on average six boutons at NMJ 6/7 in both control and *elff* LOF embryos (Figure 7A-C). Thus, Elff does not regulate initial bouton formation during embryogenesis, consistent with the observation that *elff* mutants display normal bouton number at the L3 stage (Figure 1F-H).

**Figure 7.**
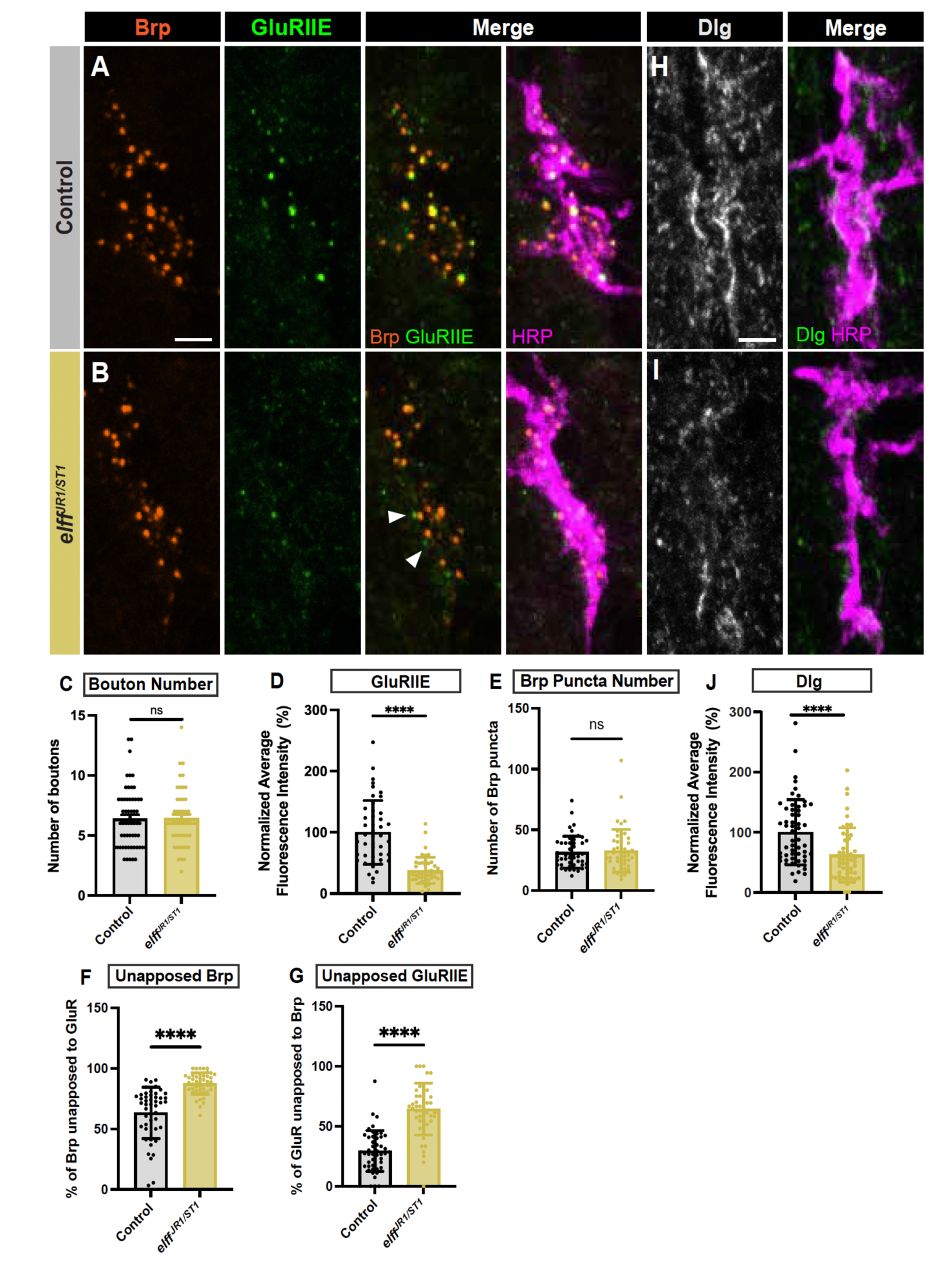
Elff promotes initial assembly of the postsynaptic compartment. (A-B) Representative Z-projections of embryonic stage 17 NMJs from the indicated genotypes stained for Brp (orange), GluRIIE (green), and HRP (magenta). Scale bar: 2 µm. (C) Quantification of bouton number per NMJ. Data are mean ± SD (control: 6.386 ± 2.505, *elff^JR1/ST1^*: 6.417 ± 2.395). (D) Quantification of GluRIIE intensity. Data are mean values normalized to control ± SD (control: 100.0 ± 52.1, *elff^JR1/ST1^*: 37.9 ± 21.4). (E) Quantification of Brp puncta number per NMJ (control: 31.51 ± 13.14, *elff^JR1/ST1^*: 32.72 ± 17.64). (F) Quantification of the percentage of Brp puncta unopposed to GluR (control: 63.35 ± 21.26, *elff^JR1/ST1^*: 87.55 ± 9.002). (G) Quantification of the percentage of GluRIIE puncta unopposed to Brp (control: 29.45 *±* 17.0, *elff^JR1/ST1^*: 64.38 ± 21.70). (H-I) Representative Z-projections of stage 17 NMJs from the indicated genotypes stained for Dlg (green) and HRP (magenta). Scale bar: 2 µm. (J) Quantification of Dlg intensity (control: 100.0 ± 54.1, *elff^JR1/ST1^*: 62.4 ± 45.3). Significance determined by Mann-Whitney test [ns, not significant, ****p<0.0001]. n ≥ 45, Animals ≥ 15.

We next assayed embryonic synapse formation using the cytomatrix protein Bruchpilot (Brp) as a presynaptic marker (Wagh et al., 2006) and GluRIIE as a postsynaptic marker. In control embryos, we find clear evidence of ongoing synapse formation with larger, more mature, presynaptic Brp puncta in close apposition to postsynaptic GluR clusters (Figure 7A). In *elff* mutants, while we do not find a difference in presynaptic Brp relative to controls, we detect a greater than 60% decrease in postsynaptic GluRIIE levels (Figure 7A-E). In addition, while in control embryos larger GluRIIE puncta are closely apposed to Brp puncta, in *elff* mutants this tight apposition is lost (arrows in Figure 7A-B). Together, these data indicate a requirement for *elff* in the formation of postsynaptic GluR complexes, and furthermore, in the critical role of aligning these complexes with presynaptic active zones for rapid neurotransmission.

Finally, we investigated whether Elff promotes postsynaptic assembly beyond the formation of GluR clusters. Since Elff is required for both alpha-Spectrin and Dlg localization at the L2 and L3 stages, we wanted to test whether this function is for initial localization of these proteins to the postsynaptic specialization, or rather for their maintenance. While we were not able to achieve consistent alpha-Spectrin labeling in controls at embryonic stage 17, we found robust Dlg expression in controls at this early stage (Figure 7H). In contrast, we observe a roughly 50% reduction in Dlg levels at embryonic *elff* mutants (Figure 7H-J). Together, these findings argue that Elff is required for multiple essential features of postsynaptic assembly.

### Elff is essential for transcellular nanoalignment of pre-and postsynaptic specializations

Our physiological analysis indicated that *elff* mutants do not exhibit reduced quantal content but do exhibit reductions in both mini amplitude and frequency (Figure 4), arguing that postsynaptic function is selectively impaired. Interestingly, our embryonic analysis revealed a defect in apposition of pre-and postsynaptic specializations in stage 17 *elff* nulls, suggesting that nanoscale synapse architecture is aberrant. To test this hypothesis at larval stage NMJs, we employed high-resolution Airyscan confocal imaging and and an Imaris analysis pipeline to quantify transsynaptic alignment in *elff* mutants using the presynaptic marker Brp and the postsynaptic marker GluRIIE. As previously described, *elff* mutant NMJs display diminished GluR levels. Thus, confocal gain was increased in these preparations to enable clear visualization of GluR clusters (see Methods).

Focusing first on the presynaptic side, we find that active zone density is unaltered in *elff* nulls, though the size of individual Brp puncta is modestly reduced (Figure 8A-B, D-E). We then investigated whether Elff is required for alignment of pre-and postsynaptic compartments at the L3 stage. In control NMJs, the overwhelming majority of individual Brp and GluR dyads are observed as overlapping puncta when viewed in single slices of Airyscan images (Figure 8A, F). In stark contrast, we find an 8-fold increase in unapposed Brp puncta in *elff* nulls (white arrowheads in Figure 8B, F) and a 7-fold increase in unapposed GluRIIE puncta in *elff* nulls (yellow arrowheads in Figure 8B, G). In particular, the presence of large GluR fields unopposed to presynaptic specializations (yellow arrowheads in 8B) in *elff* mutants argues strongly that this phenotype is not merely the result of an inability to detect GluR clusters in the mutants, but rather a combination of reduced and misaligned receptor clusters.

**Figure 8.**
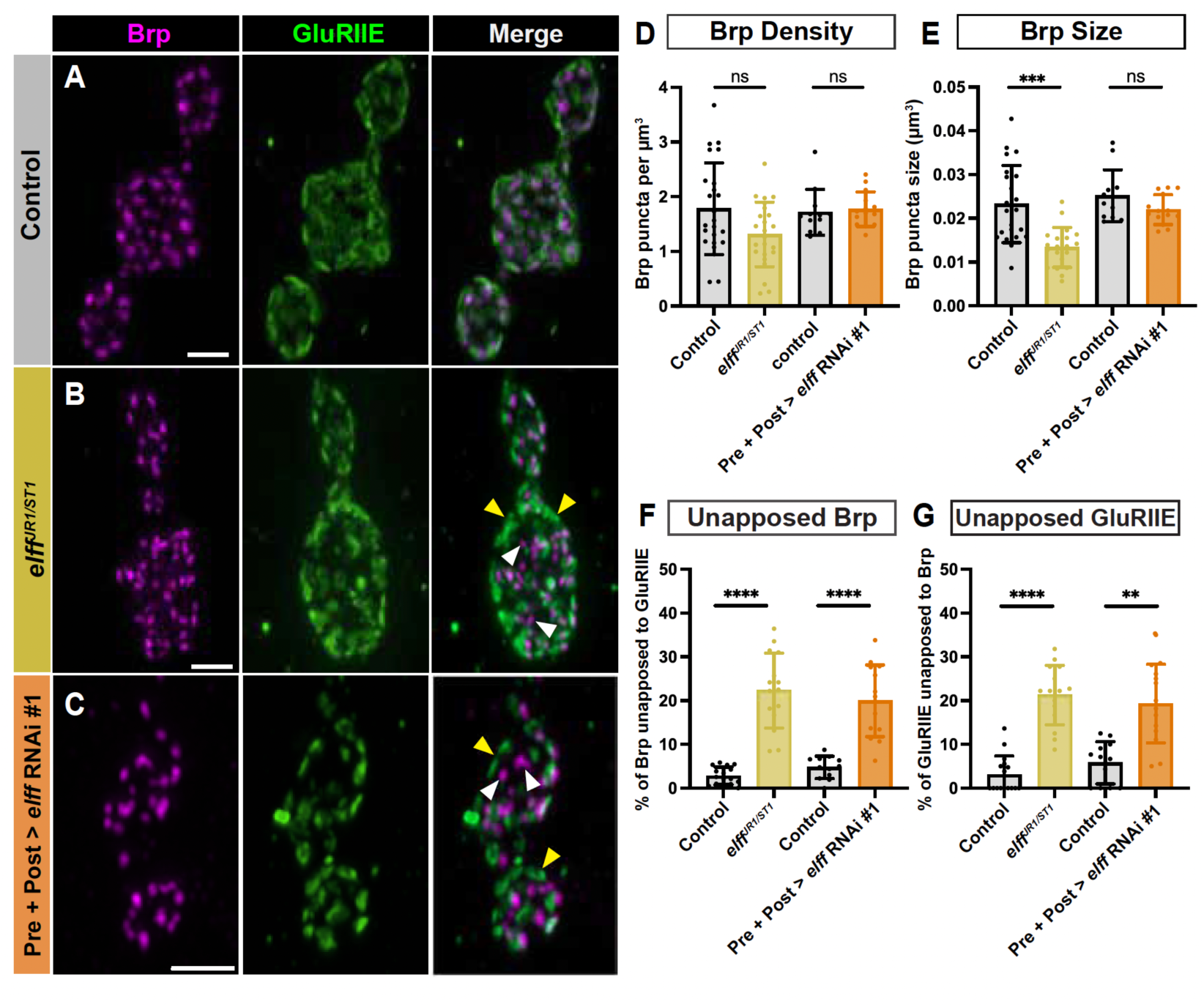
Elff is essential for transcellular nanoalignment of pre-and postsynaptic specializations. (A-C) Representative Airyscan Z-projections of boutons from the indicated genotypes stained for Brp (magenta) and GluRIIE (green). White arrowheads indicate unapposed Brp. Yellow arrowheads indicate unopposed GluRIIE. Scale bars: 2 µm. (D) Quantification of Brp density. Data are mean ± SD (control: 1.8 ± 0.8, *elff^JR1/ST1^*: 1.3 ± 0.6; control: 1.7 ± 0.4, Pre+Post > *elff* RNAi #1: 1.8 ± 0.3). (E) Quantification of Brp puncta size (control: 0.02 ± 0.01, *elff^JR1/ST1^*: 0.01 ± 0.00; control: 0.03 ± 0.01, Pre+Post > *elff* RNAi #1: 0.02 ± 0.00). (F) Quantification of the percentage of unapposed Brp (control: 2.8 ± 2.0, *elff^JR1/ST1^*: 22.3 ± 8.6; control: 4.8 ± 2.6, Pre+Post > *elff* RNAi #1: 20.0 ± 8.2). (G) Quantification of the percentage of unapposed GluRIIE (control: 3.1 ± 4.2, *elff^JR1/ST1^*: 21.3 ± 6.8; control: 5.8 ± 4.8, Pre+Post > *elff* RNAi #1: 19.3 ± 9.0). Significance determined by Kruskal-Wallis test with Dunn’s multiple comparisons (D-E, G) or one-way ANOVA with Tukey’s multiple comparisons test (F) [ns, not significant,**p<0.01, ***p<0.001, ****p<0.0001]. n ≥ 12, Animals ≥ 6.

The widespread misalignment of pre-and postsynaptic compartments in *elff* mutants argues that this extracellular synaptic adhesion molecule acts in the synaptic cleft to bring the pre-and postsynaptic compartments into register. To define the specific cellular requirement for Elff in the alignment of pre-and postsynaptic specializations, we employed cell-type specific RNAi-mediated knockdown. Similar to our GluR analysis, neither *elff* knockdown in motoneurons nor muscle results in overt defects in synapse apposition (data not shown). However, simultaneous knockdown of *elff* in both pre-and postsynaptic compartments leads to phenotypes comparable to that observed in the nulls (Figure 8A-G); with the exception of Brp size, which is not significantly affected in the RNAi condition (Figure 8E).

Specifically, combined pre-and postsynaptic *elff* knockdown results in a five-fold increase in unopposed Brp puncta (white arrowheads in Figure 8C, F) and a four-fold increase in unopposed GluR puncta (yellow arrowheads in Figure 8C, G) relative to driver-only controls. These data argue that Elff is sufficient when expressed in either the motoneuron or the muscle. Moreover, while Elff has a predicted GPI linkage, the absence of a cell-type specific requirement raises the possibility that Elff can act cell non-autonomously in this context (see Figure 9 and Discussion). The penetrance of the apposition phenotypes in *elff* null and RNAi backgrounds are at least as strong as those observed in key synapse organizers including Neuroligin and Teneurin (Banovic et al., 2010; Mosca et al., 2012), indicating that Elff is a key, novel synapse organizer at the Drosophila NMJ (Figure 9).

**Figure 9.**
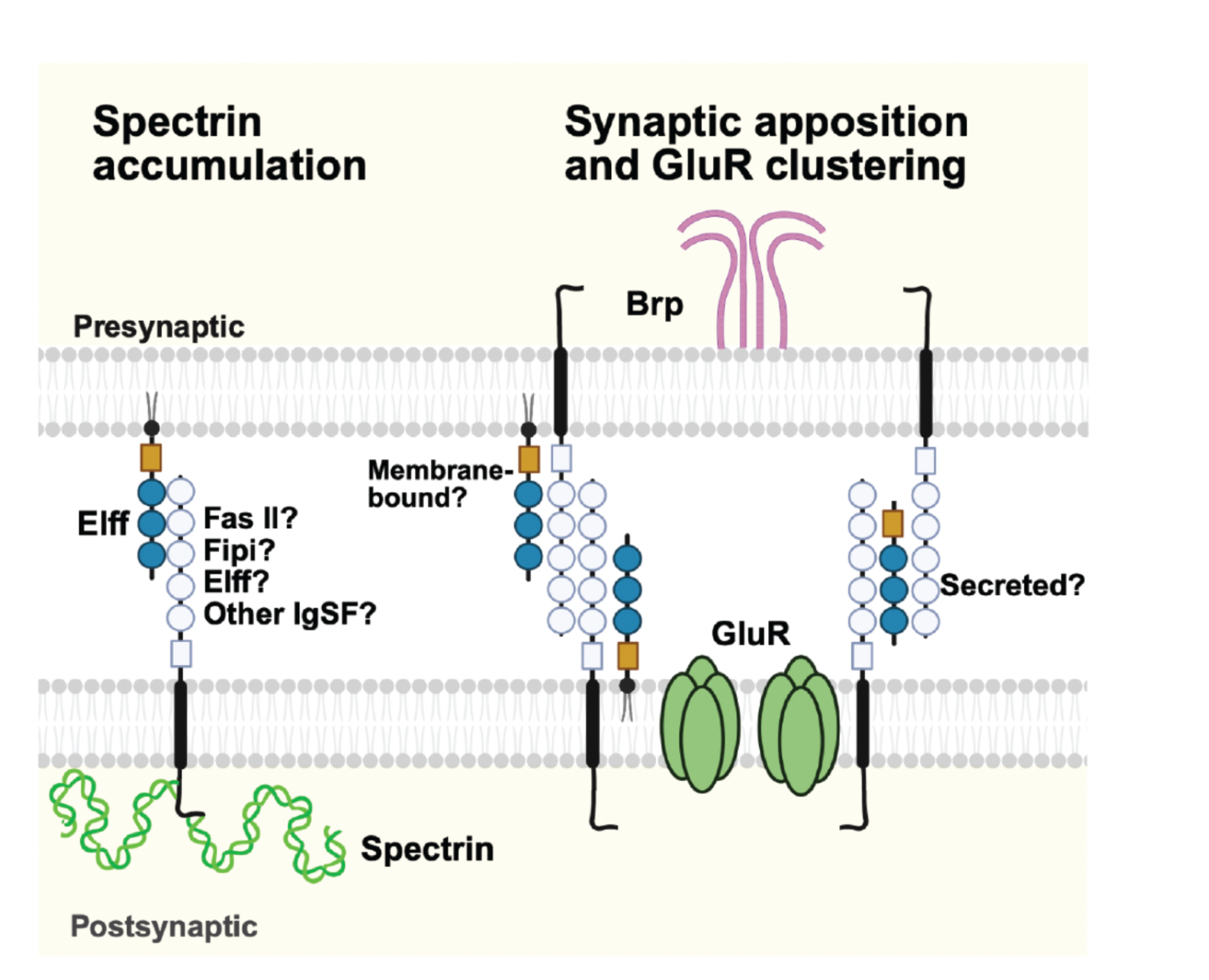
A model of distinct mechanisms for Elff’s functions in postsynaptic maturation and glutamate receptor clustering. Left: Elff is required in trans for postsynaptic accumulation of spectrin. Its postsynaptic partner(s) and other possible interactors are not known. Right: Elff facilitates glutamate receptor clustering and synaptic apposition in cis and/or in trans by stabilizing transsynaptic interactions through an unknown mechanism. In this context, Elff may act as a membrane-bound protein and/or as a secreted adaptor for other adhesion proteins in the extracellular matrix of the synaptic cleft.

## Discussion

The number of neuronal adhesion molecules expressed on growth cones and at synapses begs the question of the degree to which they have unique versus redundant functions. While unique functions for some were discovered in pioneering genetic screens for axon guidance and targeting mutants (Seeger et al., 1993; Lin et al., 1994a; Tessier-Lavigne and Goodman, 1996; Sink et al., 2001), redundancy among many others is proposed to ensure robustness of synapse formation and plasticity (Piechotta et al., 2006; Shen and Scheiffele, 2010). For example, single mutants of many synaptic adhesion molecules display only modest defects in motor axon targeting or NMJ growth in Drosophila (Nose, 2012; McLaughlin et al., 2016; Ashley and Carrillo, 2024). The prevalence of subtle phenotypes in null mutations of conserved synapse organizers raises the possibility of widespread molecular redundancy among adhesion molecules in synapse formation and maturation. In contrast to this prevailing model, here we demonstrate a clear requirement for the IgSF protein Elff in postsynaptic assembly and function in the absence of overt defects in either presynaptic function or NMJ formation and expansion. We discuss below our findings in the context of transsynaptic nanoalignment, postsynaptic assembly and maturation (Figure 9).

### Elff serves specific functions in subsynaptic organization

As a putative Fas II binding partner expressed in the developing and adult CNS, we hypothesized that Elff would be required for organismal viability and NMJ expansion. This hypothesis was particularly attractive since Elff is selectively expressed in the α_1_β_1_ and γ neurons of the adult mushroom body akin to Trio and Crimpy, key components of the widely-studied BMP signaling pathway at the Drosophila NMJ (Ball et al., 2010; James et al., 2014; Croset et al., 2018). However, *elff* nulls are viable and do not display alterations in gross morphology of the NMJ. Moreover, Fas II levels at the NMJ are unchanged in *elff* nulls relative to controls, arguing that Elff is not required for Fas II expression.

While Elff is not required for NMJ growth, it is critical for postsynaptic assembly and maturation. Specifically, both essential and alternative GluR subunit abundance is dramatically reduced at *elff* mutant NMJs. To determine if this reflects a failure of postsynaptic assembly or maintenance, we analyzed embryonic and L2 stage NMJs. We drew two main conclusions from these studies. First, our embryonic analysis argues that Elff contributes to initial assembly of the postsynaptic compartment since we find defects from embryonic late stage 17 onward, which is the earliest we observe apposed Brp-GluR puncta in controls. Second, our L2 analysis indicates a more stringent requirement for Elff at the L2 versus L3 stage, e.g., Dlg and GluRIIE are virtually undetectable at L2, while at L3, they are reduced but still visible. These findings are consistent with many others demonstrating the existence of homeostatic pathways at the NMJ (Dickman and Davis, 2009; Davis, 2013; Perry et al., 2017; Goel and Dickman, 2018; Goel et al., 2019), and argue that in the absence of Elff, alternative molecular mechanism(s) compensate for its absence. More broadly, the relative strength of the *elff* L2 phenotype relative to L3 suggests that the strength of mutant phenotypes related to synapse organization (as opposed to bouton number) may be diminished by phenotypic analyses focused solely on L3 animals.

The viability and only modestly impaired motility of homozygous (*elff^JR1/JR1^*and *elff^ST1/ST1^*) and transheterozgous mutants (*elff^JR1/ST1^*) was initially surprising to us given the dramatic loss of GluR clusters. In fact, our data are consistent with studies from multiple groups indicating that even 5% of normal GluR cluster abundance is sufficient to support larval motility and viability due to the presence of a safety factor to ensure muscle contraction (Marrus et al., 2004; Qin et al., 2005; Schmid et al., 2006; Kim et al., 2012). In addition, our finding that aspects of presynaptic organization and function including active zone density and quantal content are normal in *elff* mutants is consistent with previous work establishing that total, or near-total, GluR loss does not disrupt presynaptic organization (Marrus et al., 2004; Schmid et al., 2006; Kim et al., 2012).

Elff is necessary for pre-and postsynaptic apposition, glutamate receptor localization, and postsynaptic protein expression. This set of *elff* phenotypes at the NMJ raises the question about their possible interdependency. GluR field size is not reduced in the absence of either Dlg (Chen and Featherstone, 2005) or alpha-Spectrin (Pielage et al., 2005), arguing that their loss in *elff* mutants is not responsible for the GluR phenotype. In contrast, both Dlg levels and SSR thickness are reduced in mutants of essential GluR subunits or the auxiliary subunit Neto (Schmid et al., 2006; Kim et al., 2012). These prior findings, coupled with the identity of Elff as a secreted/GPI-linked IgSF, are consistent with the hypothesis that Elff is required (directly or indirectly) for GluR localization, and in its absence, GluRs are not properly localized, leading to Dlg loss. In contrast, we propose that Elff controls alpha-Spectrin independently of GluRs and Dlg since unlike GluRs and Dlg, Elff acts exclusively in trans to promote alpha-Spectrin levels (Figure 9).

Using high-resolution Airyscan confocal imaging, we found extensive defects in pre-and postsynaptic apposition (Figure 8); indeed, loss of Elff results in unapposed Brp puncta and GluR clusters with a frequency similar to that observed in mutants of long-studied transsynaptic adhesion proteins including Neuroligin and the Teneurins (Banovic et al., 2010; Mosca et al., 2012). Is the loss of GluRs in *elff* mutants also likely to underlie the breakdown of apposition? Loss of GluRs at the Drosophila NMJ does not lead to extensive defects in apposition of active zones to postsynaptic densities (Schmid et al., 2006; Kim et al., 2012). For example, strong hypomorphic alleles of the auxiliary GluR subunit Neto result in near total loss of GluR clusters without causing significant apposition defects (Kim et al., 2012). Thus, we propose that Elff’s function in trans-synaptic nanoalignment is distinct from its function localizing GluRs and suggests that Elff regulates adhesive nanodomains required for the precise registration of pre-and postsynaptic specializations (Biederer et al., 2017). Identification of extracellular Elff binding partners at the NMJ is essential to elucidate Elff-dependent signaling mechanisms.

We undertook RNAi experiments to validate phenotypes observed in the *elff* nulls and to address cell-type specific requirements for Elff. Simultaneous knockdown of *elff* pre-and postsynaptically leads to defects in apposition and GluR localization comparable to that observed in our null alleles (Figures 5 and 8), while *elff* knockdown in either the pre-or postsynaptic cell does not. These findings are consistent with the identity of Elff as a secreted/GPI-linked protein deposited in the synaptic cleft from both pre-and postsynaptic cells. These results are distinct from the requirements of Elff in the regulation of alpha-Spectrin and Dlg, consistent with the model that Elff is a member of distinct protein complexes serving different functions at the synapse (Figure 9).

Our functional studies are in line with Elff expression data (Figure 1). An *elff* MIMIC protein trap indicates broad neuronal expression in the third-instar larval CNS and the adult brain. We employed CRISPR-mediated genome engineering to generate an endogenous N-term epitope tagged protein, Elff^sfGFP-N^, which is expressed strongly at presynaptic NMJ terminals and at lower levels in postsynaptic muscle. While Elff localizes to the NMJ, it does not appear overtly punctate. Thus, although Elff is essential for GluR localization and alignment to active zones, our data do not indicate that Elff is itself localized to active zones or GluR clusters. One caveat is that Elff^sfGFP-N^ is an allele indicating that the GFP tag compromises Elff function. Future analysis of Elff localization at the NMJ necessitates generating additional tools for protein visualization.

### Identifying Elff-mediated signaling pathways

To our knowledge, the specific complement of NMJ phenotypes observed in *elff* mutants is unique among Drosophila mutants. Specifically, the combination of defects in apposition and postsynaptic maturation, alongside normal NMJ growth, argues that Elff does not influence the activity of key regulators of synapse adhesion including Nrx, Nlg, and the BMP pathway, since all regulate baseline NMJ growth (Marqués et al., 2002; Li et al., 2007; Banovic et al., 2010; Hoover et al., 2019). Might Elff homologs provide insight into the identity of Elff-mediated signaling pathways? Structurally, Elff is closely related to the *C. elegans* IgSF protein Rig-3, which regulates Acetylcholine receptor levels via inhibiting the Ror receptor tyrosine kinase family member CAM-1 and the Wnt pathway (Babu et al., 2011). In Drosophila, loss of Wnt pathway components results in phenotypes overlapping that of *elff* nulls (Packard et al., 2002; Restrepo et al., 2022), suggesting a possible relationship between Wnt signaling and Elff. In a similar vein, it will be important to probe possible functional relationships among Elff and the three Drosophila Ror family members (Nrk, Ror, and Drl-2).

Elff shares a close structural relationship with the Drosophila protein Fipi, and both proteins were identified as extracellular binding partners of Fasciclin II in a systematic cataloging of pairwise extracellular interactions among LRR and IgSF proteins in Drosophila (Özkan et al., 2013). Fipi’s molecular and cellular function in the CNS has not been reported, though a behavioral phenotype has been reported in male courtship song (Fedotov et al., 2018). Recently, Fipi and Elff have been shown to act in parallel in the developing wing disk to restrain EGFR activity. In this capacity, they are proposed to act both independently and in conjunction with Fas II (Garcia-Alonso, 2024). We do not currently favor the model that Fipi functions analogously to Elff at the NMJ since neither RNAi-mediated knockdown nor overexpression of Fipi display phenotypes akin to that described in this work (MNW and HTB, unpublished data). Future work will be required to determine whether Fipi contributes to NMJ development or function.

Finally, it will be critical to test if Elff regulates aspects of Fas II-dependent signaling at the NMJ. Elff is highly unlikely to be an obligate partner of Fas II since loss of Fas II results in early larval lethality, and mutant NMJs exhibit striking growth defects (Schuster et al., 1996b, 1996a; Neuert et al., 2019).

Consistent with the phenotypic differences between *elff* and *fas II* mutants, we do not detect a change in overall Fas II levels at the NMJ in *elff* nulls (Figure S1). Notably, in addition to the well-known roles of Fas II in growth regulation and adhesion, it promotes postsynaptic specification during embryonic NMJ development (Kohsaka et al., 2007). Indeed, loss of Fas II results in phenotypes overlapping those observed with loss of Elff, raising the possibility that Elff sculpts Fas II function. Moving forward, defining Elff-dependent signaling mechanisms will elucidate the molecular pathways driving transsynaptic nanoarchitecture and postsynaptic assembly at the NMJ.

## Methods

### Drosophila stocks

Flies were reared on standard medium in incubators set to 29°C (for Gal4-UAS experiments) or 25°C (all other experiments). Males and females were used for all experiments. The following fly lines were used: iso31 (gift of Masashi Tabuchi, Case Western Reserve University), w^1118^, D42Gal4, 24BGal4, ElavGal4, Elff^MI02791-GFSTF.1^ (BDSC #60531), UAS-*elff* RNAi (BDSC #64879, referred in this work as *elff* RNAi #1), UAS-*elff* RNAi (VDRC #40821, referred in this work as *elff* RNAi #2), vasa>Cas9 (BDSC#51324), and piggyBac transposase (BDSC #8283). Lines generated for this study include *elff^JR1^*, *elff^ST1^*, *elff^sfGFP-N^*, and UAS-Elff (see below). For Gal4-UAS experiments, progeny of respective driver lines crossed to iso31 served as controls. For all other experiments, iso31 flies (Figures 1, 2, 3, 5) or progeny of iso31 flies outcrossed to w^1118^ (Figures 4, 6, 7, 8) were used as controls.

### Generation of alleles

The *elff^JR1^* and *elff^ST1^* alleles were generated via published CRISPR/Cas9 genome engineering strategies (Gratz et al., 2013, 2015). Briefly, for *elff^JR1^*, ∼1kb targeting homology arms flanking the coding region of *elff* were synthesized and subcloned (GenScript, Inc.) into the pHD-DsRed-attP-w+ plasmid (Addgene 80898) to generate a donor template for homology-directed repair. A CG33543-targeting guide RNA (gRNA) was generated by inserting a target sequence (5’-TTGCTGGCGATGGGGCAACA-3’) into the pU6-3-gRNA plasmid (DGRC 1362) via site-directed mutagenesis (NEB Cat #E0554S). The homology-directed repair and gRNA plasmids were then injected into vasa>Cas9 embryos (BDSC 51324; BestGene, Inc.) to create CG33543 deletions, in which 9.4kb of the endogenous CG33543 gene (2Lcomplement:1996930..2006339) were replaced with an attP-loxP-DsRed-loxP cassette.

### Generation of UAS lines

CG33543 cDNA was synthesized and subcloned into the pACU UAS transformation vector (Chun Han, Addgene 58373; GenScript, Inc.). This donor vector was then injected into embryos of an attP transposable element insertion stock (BDSC 36304; BestGene, Inc.) to generate a stable Drosophila line expressing CG33543 cDNA downstream of UAS regulatory regions.

### Immunohistochemistry

The following commercially available primary antibodies were used: anti-Bruchpilot (DSHB, nc82) [1:100]; anti-Discs large (DSHB, 4F3) [1:500 third instar, 1:100 second instar, 1:100 embryo]; anti-alpha Spectrin (DSHB, 3A9) [1:100 third instar, 1:50 second instar]. Primary antibodies for GluR subunits were provided by Dion Dickman (University of Southern California): guinea pig anti-GluRIIA [1:1000]; rabbit anti-GluRIIB [1:1000]; rabbit anti-GluRIIC [1:2500], rabbit anti-GluRIIE [1:2000 third instar, 1:500 second instar, 1:500 embryo]. Secondary antibodies used in this work were goat anti-mouse Alexa Fluor 488-, 568-, and 647-conjugated IgG (Invitrogen, A-11029; A-11031; A-32728), goat anti-rabbit Alexa Fluor 488-conjugated IgG (Invitrogen, A-11034), and goat anti-guinea pig Alexa Fluor 488-conjugated IgG (Invitrogen, A-11073). All secondary antibodies were used at a concentration of 1:300. HRP staining was performed by incubating samples with Alexa Fluor 647-conjugated or DyLight 594-conjugated anti-HRP (Jackson ImmunoResearch 123-585-021, 123-605-021) at 1:200 alongside secondary antibodies. GFP staining was performed by incubating samples with GFP Polyclonal Antibody, Alexa Fluor 488 (Invitrogen, A-21311) [1:500 third instar, 1:400 adult brain].

### Adult brain dissections

Adult flies were anesthetized on ice and decapitated. Brains were then rapidly dissected from the heads in ice-cold PBS and fixed for 20 min with 4% PFA in PTX10 (1% Triton X-100 in PBS). Fixed brains were washed with PTX10 and incubated for 2 hr in blocking buffer (5% normal goat serum, 0.3% Triton X-100 in PBS), followed by an overnight incubation at 4°C with conjugated GFP antibody diluted in PTX (0.1% Triton X-100 in PBS). The following day, brains were washed 3×30 min in PTX10, then placed in SlowFade Gold mountant (Invitrogen S36936) for approximately 30 min (stationary). Brains were then mounted as previously described (Kelly et al., 2017). Unless otherwise noted, all fixation, incubation, and washing steps were performed at room temperature (RT) while nutating.

### Larval NMJ dissections

Second-and third-instar larvae were dissected in PBS and fixed for 10 min in Bouin’s fixative. Larval fillets were then blocked at RT in either 1% (third instar) or 5% (second instar) NGS in PTX and incubated overnight with primary antibodies at 4°C. Fillets were then washed in 1% NGS and incubated with secondary antibodies for 2-4 hours at RT. Following a final wash in PTX, fillets were mounted in Pro-Long Gold (Thermo Fisher #P36930).

### Embryo dissections

Overnight collections of Drosophila embryos were dechorionated in 50% bleach for 4 minutes. After dechorionation, stage 17 embryos were selected, fixed in 2 ml methanol and 2 ml heptane with gentle rocking for 2 minutes, and devitellinized by 1 minute of shaking in 6 ml methanol. Devitellinized embryos were partially dissected in a droplet of PBS on a glass microscope slide by making a small opening in the ventral cuticle, followed by a second fixation in cold methanol for 5 minutes. Fully fixed embryos were blocked in 5% NGS, then incubated with primary antibodies and conjugated HRP antibody overnight at 4°C and an additional 4 hr at RT. Embryos were then washed in 5% NGS for 3 washes of 5 minutes and incubated with secondary antibodies overnight at 4°C. After staining, embryos were fully filletted on a microscope slide in 50% glycerol and mounted in Pro-Long Gold.

### Confocal microscopy

Standard confocal images were acquired with a Zeiss LSM 800 laser scanning microscope using a 20x objective for adult brains, a 40x objective for third-instar larvae, a 63x objective for second-instar larvae, and a 100x objective for stage 17 embryos. The Zeiss Airyscan software was used with a 63x objective for obtaining images in Figure 8 only. For intensity quantification, laser intensity and gain were maintained across an experiment. For nascent bouton quantification, laser intensity and gain were optimized for each Z-stack.

### Electrophysiology

Electrophysiological recordings were performed as previously described (Zhang et al., 2010) with minor modifications. Male third-instar larvae were dissected in low-Ca²⁺ hemolymph-like saline (HL3; in mM: 70 NaCl, 5 KCl, 10 MgCl₂, 10 NaHCO₃, 115 sucrose, 5 trehalose, 5 HEPES, pH 7.2)(Zhang and Stewart, 2010) containing 0.2 mM Ca²⁺. Sharp intracellular recordings were performed in HL3 saline containing 0.6 mM Ca²⁺ from muscle 6 of abdominal segments A3 and A4 using borosilicate electrodes (resistance 10–25 MΩ) filled with 3 M KCl. Recordings were conducted using a Nikon FN1 microscope with a 40X/0.80 NA water-dipping objective, and acquired with an AxoClamp 900A amplifier, Digidata 1550B digitizer, and pClamp 11 software (Molecular Devices). Signals were digitized at 10 kHz and filtered at 1 kHz. Miniature excitatory junctional potentials (mEJPs) were recorded in the absence of stimulation and cut motor axons were stimulated to elicit evoked EJPs. For each cell, at least 60 mEJPs were analyzed using Mini Analysis (Synaptosoft) to calculate mean mEJP amplitude. EJPs were elicited using an Isolated Pulse Stimulator (Model 2100, A-M Systems), at a frequency of 0.2 Hz with intensity adjusted per cell to consistently activate compound responses from both type Ib and Is motor neurons innervating muscle 6. At least 30 consecutive EJPs were recorded and analyzed in Clampfit to obtain mean EJP amplitude. Quantal content was estimated by dividing mean EJP amplitude by mean mEJP amplitude. Muscle input resistance (R in) and resting membrane potential (V rest) were monitored throughout. Recordings were included in the analysis only if V rest was between –50 and –80 mV and R in was ≥5 MΩ.

### Larval Crawling

Prior to behavioral testing, 100mm x 20mm Petri dishes were prepared with a layer of 1% agarose dyed with 0.6% brilliant black fountain ink (Pelikan). On testing day, three third-instar larvae were transferred to the center of a black-dyed agarose plate containing a thin layer of dH_2_O for moisture. Larvae were recorded for 5 minutes using a 48MP phone camera. All movies were then converted to 0.2 frames per second in AVI format and uploaded to ImageJ (v2.16.0). Using the Manual Tracking plugin, larval position was indicated every 5 s (60 frames total) to measure total distance crawled and average velocity. Independent experimental trials were performed around the same time of day.

### Image Analysis

Bouton Number: Bouton numbers were manually counted from muscle 4 on segments A2-A5 in third instar larvae, from muscle 6/7 on segments A3-A4 in second instar larvae, and from muscle 6/7 on segments A2-A5 in embryos.

Nascent Boutons: NMJs were manually scored from muscles 6/7, segment A2. Nascent boutons were quantified as those having HRP signal with no appreciable Dlg signal present.

Fluorescence Intensity, Puncta Measurements, and Apposition: Quantification of fluorescence intensity, puncta density/size, and apposition was done using Imaris 9.7.1 (Bitplane). For third instar NMJs, intensity measurements of synaptic alpha-Spectrin and Dlg were made by generating Surfaces from the respective channels. For fluorescence intensity measurements of GluR and Fas II, the Surfaces function was used on the HRP channel to outline neuronal membranes. In each case, the resulting surface was manually processed by an experimenter to isolate the NMJ being analyzed, and the mean fluorescence intensity value of the channel of interest within the generated surface was captured. To quantify Brp puncta density and size in third instar NMJs, the Surfaces function was used on the HRP channel and manually processed to isolate the NMJ. The resulting HRP surface was then used to mask Brp signal within the NMJ. Using the Spots function, Brp puncta within the masked channel were identified and quantified under the following parameters: algorithm set to different Spot sizes (region growing), background subtraction enabled, estimated diameter of 0.2 µm, and Spots region from absolute intensity with automatic region thresholding enabled. To calculate puncta density, the number of puncta per NMJ was divided by the volume of the HRP surface. For embryonic NMJs, Brp puncta number and Dlg intensity were quantified using slightly modified versions of the third instar pipelines, while GluR intensity was measured by identifying GluR puncta via a Spots function on GluR signal within the masked HRP surface and capturing mean fluorescence intensity within those spots. To quantify apposition of Brp and GluRIIE in both third instar and embryonic NMJs, Brp spots and GluRIIE clusters/spots not co-localized with their respective synaptic partner marker were manually counted and divided by the total number of Brp spots or GluRIIE clusters, respectively, at each NMJ.

### Statistics

Statistical analyses were performed with Prism 10.2.1 (GraphPad). All data is from at least two independent experiments. For microscopy experiments, each data point represents one neuromuscular junction, unless otherwise noted. For electrophysiology and behavior experiments, each data point represents recordings from one animal. Shapiro-Wilks normality testing was performed on all data sets. On those demonstrating a normal distribution, an unpaired t-test for two groups or a one-way ANOVA followed by Tukey’s multiple comparisons test for three or more groups were performed. For non-normal distributions, nonparametric tests were used: a Mann-Whitney test for comparisons of two groups or a Kruskal-Wallis followed by Dunn’s multiple comparisons test for three or more groups. For categorical data, a chi-square test was used. p-values, statistical tests, ‘n’, and number of animals used are reported in each figure legend.

## Acknowledgements

We are indebted to Flybase for their work annotating the *elff* locus. We thank the Bloomington Drosophila Stock Center for fly stocks and the Developmental Studies Hybridoma Bank for antibodies. We are grateful to Dion Dickman for anti-GluR antibodies and Masashi Tabuchi for stocks. We thank members of the Broihier lab for helpful discussions and comments on the manuscript. This work was supported by the NIH under the awards R01NS095895 to HTB and R01NS078179 to KOCG.

---

**Supplemental Figure S1.**
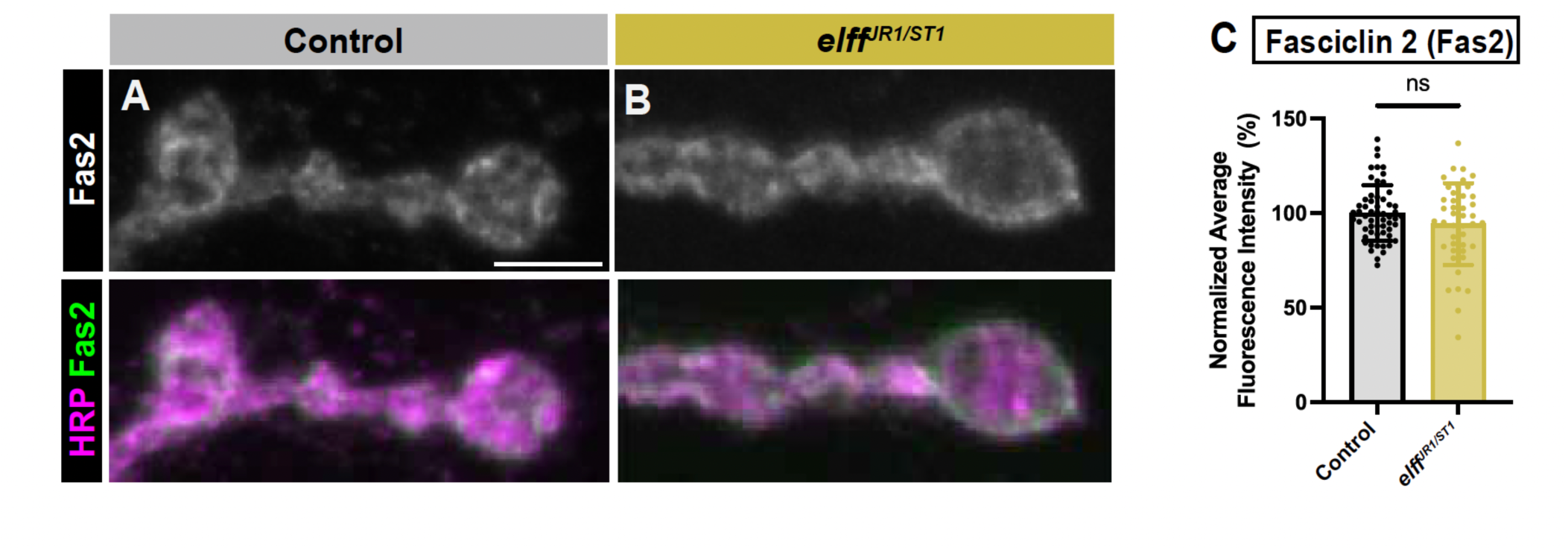
(A-B) Z-projections of NMJ boutons of the indicated genotypes stained for Fas2 (green) and HRP (magenta). (C) Quantification of Fas2 fluorescence intensity. Data are mean values normalized to control ± SD (control: 100.0 ± 14.71, *elff^JR1/ST1^*: 94.19 ± 21.59). Significance determined by unpaired t-test [ns, not significant]. n ≥ 43, Animals ≥ 11.

**Supplemental Figure S2.**
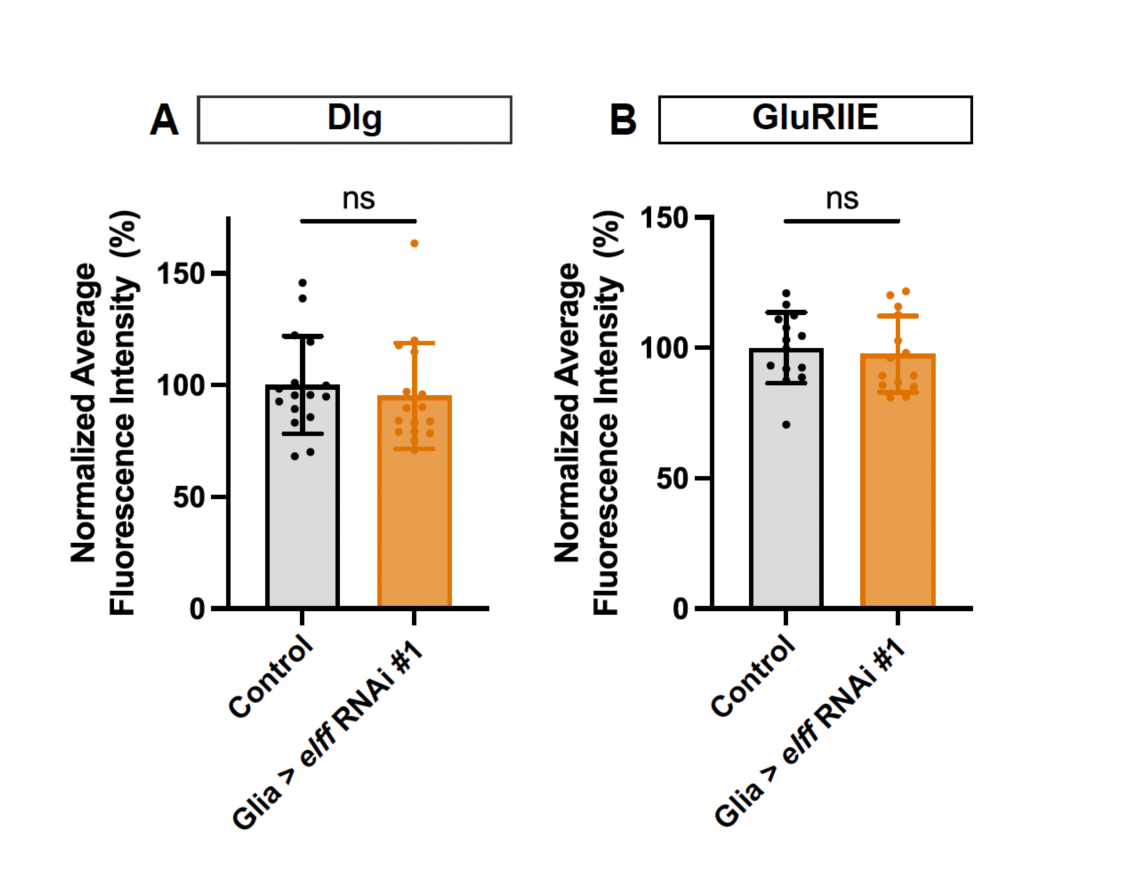
(A-B) Quantification of fluorescence intensity of Dlg (A) and GluRIIE (B) staining in third instar larval NMJs at muscle 4 from the indicated genotypes. The Gal4 driver used for glia is RepoGal4. Data are mean values normalized to control ± SD for both Dlg (control: 100.0 ± 21.78, glia > *elff* RNAi: 95.15 ± 23.68) and GluRIIE (control: 100.0 ± 13.60, glia > *elff* RNAi: 97.54 ± 14.62). Significance determined by Mann-Whitney test (A) or unpaired t-test (B) [ns, not significant]. n ≥ 14, Animals ≥ 4.

## References

Aonurm-Helm A, Jaako K, Jürgenson M, Zharkovsky A (2016) Pharmacological approach for targeting dysfunctional brain plasticity: Focus on neural cell adhesion molecule (NCAM). Pharmacol Res 113:731–738.

Ashley J, Carrillo RA (2024) The Drosophila Larval Neuromuscular Junction: Developmental Overview. Cold Spring Harb Protoc 2025:pdb.top108449.

Ashley J, Packard M, Ataman B, Budnik V (2005) Fasciclin II Signals New Synapse Formation through Amyloid Precursor Protein and the Scaffolding Protein dX11/Mint. J Neurosci 25:5943–5955.

Ataman B, Ashley J, Gorczyca M, Ramachandran P, Fouquet W, Sigrist SJ, Budnik V (2008) Rapid Activity-Dependent Modifications in Synaptic Structure and Function Require Bidirectional Wnt Signaling. Neuron 57:705–718.

Atwood HL, Govind CK, Wu C -F. (1993) Differential ultrastructure of synaptic terminals on ventral longitudinal abdominal muscles in Drosophila larvae. J Neurobiol 24:1008–1024.

Babu K, Hu Z, Chien S-C, Garriga G, Kaplan JM (2011) The Immunoglobulin Super Family Protein RIG-3 Prevents Synaptic Potentiation and Regulates Wnt Signaling. Neuron 71:103–116.

Ball RW, Warren-Paquin M, Tsurudome K, Liao EH, Elazzouzi F, Cavanagh C, An B-S, Wang T-T, White JH, Haghighi AP (2010) Retrograde BMP Signaling Controls Synaptic Growth at the NMJ by Regulating Trio Expression in Motor Neurons. Neuron 66:536–549.

Banovic D, Khorramshahi O, Owald D, Wichmann C, Riedt T, Fouquet W, Tian R, Sigrist SJ, Aberle H (2010) Drosophila Neuroligin 1 Promotes Growth and Postsynaptic Differentiation at Glutamatergic Neuromuscular Junctions. Neuron 66:724–738.

Biederer T, Kaeser PS, Blanpied TA (2017) Transcellular Nanoalignment of Synaptic Function. Neuron 96:680–696.

Butler SJ, Ray S, Hiromi Y (1997) klingon, a novel member of the Drosophila immunoglobulin superfamily, is required for the development of the R7 photoreceptor neuron. Development 124:781–792.

Chen K, Featherstone DE (2005) Discs-large (DLG) is clustered by presynaptic innervation and regulates postsynaptic glutamate receptor subunit composition in Drosophila. BMC Biol 3:1.

Chen K, Gracheva EO, Yu S-C, Sheng Q, Richmond J, Featherstone DE (2010) Neurexin in Embryonic Drosophila Neuromuscular Junctions. PLoS ONE 5:e11115.

Croset V, Treiber CD, Waddell S (2018) Cellular diversity in the Drosophila midbrain revealed by single-cell transcriptomics. Elife 7:e34550.

Cunningham BA, Hemperly JJ, Murray BA, Prediger EA, Brackenbury R, Edelman GM (1987) Neural Cell Adhesion Molecule: Structure, Immunoglobulin-Like Domains, Cell Surface Modulation, and Alternative RNA Splicing. Science 236:799–806.

Davis GW (2013) Homeostatic Signaling and the Stabilization of Neural Function. Neuron 80:718–728.

Davis GW, Schuster CM, Goodman CS (1997) Genetic Analysis of the Mechanisms Controlling Target Selection: Target-Derived Fasciclin II Regulates the Pattern of Synapse Formation. Neuron 19:561–573.

DePew AT, Aimino MA, Mosca TJ (2019) The Tenets of Teneurin: Conserved Mechanisms Regulate Diverse Developmental Processes in the Drosophila Nervous System. Front Neurosci-switz 13:27.

DiAntonio A, Petersen SA, Heckmann M, Goodman CS (1999) Glutamate Receptor Expression Regulates Quantal Size and Quantal Content at the Drosophila Neuromuscular Junction. J Neurosci 19:3023–3032.

Dickman DK, Davis GW (2009) The Schizophrenia Susceptibility Gene dysbindin Controls Synaptic Homeostasis. Science 326:1127–1130.

Duncan BW, Murphy KE, Maness PF (2021) Molecular Mechanisms of L1 and NCAM Adhesion Molecules in Synaptic Pruning, Plasticity, and Stabilization. Frontiers Cell Dev Biology 9:625340.

Featherstone DE, Rushton E, Rohrbough J, Liebl F, Karr J, Sheng Q, Rodesch CK, Broadie K (2005) An Essential Drosophila Glutamate Receptor Subunit That Functions in Both Central Neuropil and Neuromuscular Junction. J Neurosci 25:3199–3208.

Fedotov SA, Bragina JV, Besedina NG, Danilenkova LV, Kamysheva EA, Kamyshev NG (2018) Gene CG15630 (fipi) is involved in regulation of the interpulse interval in Drosophila courtship song. J Neurogenet 32:15–26.

Fedotov SA, Bragina JV, Besedina NG, Danilenkova LV, Kamysheva EA, Panova AA, Kamyshev NG (2014) The effect of neurospecific knockdown of candidate genes for locomotor behavior and sound production in Drosophila melanogaster. Fly 8:176–187.

Garcia-Alonso L (2024) Fasciclin 2 functions as an expression-level switch on EGFR to control organ shape and size in Drosophila. PLOS ONE 19:e0309891.

Goel P, Dickman D (2018) Distinct homeostatic modulations stabilize reduced postsynaptic receptivity in response to presynaptic DLK signaling. Nat Commun 9:1856.

Goel P, Khan M, Howard S, Kim G, Kiragasi B, Kikuma K, Dickman D (2019) A Screen for Synaptic Growth Mutants Reveals Mechanisms That Stabilize Synaptic Strength. J Neurosci 39:4051–4065.

Gratz SJ, Cummings AM, Nguyen JN, Hamm DC, Donohue LK, Harrison MM, Wildonger J, O’Connor-Giles KM (2013) Genome engineering of Drosophila with the CRISPR RNA-guided Cas9 nuclease. Genetics 194:1029–1035.

Gratz SJ, Rubinstein CD, Harrison MM, Wildonger J, O’Connor-Giles KM (2015) CRISPR-Cas9 Genome Editing in Drosophila. Curr Protoc Mol Biology 111:31.2.1-31.2.20.

Graveley BR et al. (2011) The developmental transcriptome of Drosophila melanogaster. Nature 471:473–479.

Hata K, Maeno-Hikichi Y, Yumoto N, Burden SJ, Landmesser LT (2018) Distinct Roles of Different Presynaptic and Postsynaptic NCAM Isoforms in Early Motoneuron–Myotube Interactions Required for Functional Synapse Formation. J Neurosci 38:498–510.

Hata K, Polo-Parada L, Landmesser LT (2007) Selective Targeting of Different Neural Cell Adhesion Molecule Isoforms during Motoneuron–Myotube Synapse Formation in Culture and the Switch from an Immature to Mature Form of Synaptic Vesicle Cycling. J Neurosci 27:14481–14493.

Hoover KM, Gratz SJ, Qi N, Herrmann KA, Liu Y, Perry-Richardson JJ, Vanderzalm PJ, O’Connor-Giles KM, Broihier HT (2019) The calcium channel subunit α2δ-3 organizes synapses via an activity-dependent and autocrine BMP signaling pathway. Nat Commun 10:5575.

James RE, Hoover KM, Bulgari D, McLaughlin CN, Wilson CG, Wharton KA, Levitan ES, Broihier HT (2014) Crimpy Enables Discrimination of Presynaptic and Postsynaptic Pools of a BMP at the Drosophila Neuromuscular Junction. Dev Cell 31:586–598.

Kelly SM, Elchert A, Kahl M (2017) Dissection and Immunofluorescent Staining of Mushroom Body and Photoreceptor Neurons in Adult Drosophila melanogaster Brains. J Vis Exp: JoVE:56174.

Kim Y-J, Bao H, Bonanno L, Zhang B, Serpe M (2012) Drosophila Neto is essential for clustering glutamate receptors at the neuromuscular junction. Genes Dev 26:974–987.

Kohsaka H, Takasu E, Nose A (2007) In vivo induction of postsynaptic molecular assembly by the cell adhesion molecule Fasciclin2. J Cell Biol 179:1289–1300.

Li J, Ashley J, Budnik V, Bhat MA (2007) Crucial Role of Drosophila Neurexin in Proper Active Zone Apposition to Postsynaptic Densities, Synaptic Growth, and Synaptic Transmission. Neuron 55:741–755.

Lin DM, Fetter RD, Kopczynski C, Grenningloh G, Goodman CS (1994a) Genetic analysis of Fasciclin II in drosophila: Defasciculation, refasciculation, and altered fasciculation. Neuron 13:1055–1069.

Lin DM, Fetter RD, Kopczynski C, Grenningloh G, Goodman CS (1994b) Genetic analysis of Fasciclin II in drosophila: Defasciculation, refasciculation, and altered fasciculation. Neuron 13:1055–1069.

Malchiodi-Albedi F, Ceccarini M, Winkelmann JC, Morrow JS, Petrucci TC (1993) The 270 kda splice variant of erythrocyte β-spectrin (βi∑2) segregates in vivo and in vitro to specific domains of cerebellar neurons. J Cell Sci 106:67–78.

Maness PF, Schachner M (2007) Neural recognition molecules of the immunoglobulin superfamily: signaling transducers of axon guidance and neuronal migration. Nat Neurosci 10:19–26.

Marqués G, Bao H, Haerry TE, Shimell MJ, Duchek P, Zhang B, O’Connor MB (2002) The Drosophila BMP Type II Receptor Wishful Thinking Regulates Neuromuscular Synapse Morphology and Function. Neuron 33:529–543.

Marrus SB, Portman SL, Allen MJ, Moffat KG, DiAntonio A (2004) Differential Localization of Glutamate Receptor Subunits at the Drosophila Neuromuscular Junction. J Neurosci 24:1406–1415.

McLaughlin CN, Nechipurenko IV, Liu N, Broihier HT (2016) A Toll receptor–FoxO pathway represses Pavarotti/MKLP1 to promote microtubule dynamics in motoneurons. J Cell Biology 214:459–474.

Menon KP, Carrillo RA, Zinn K (2013) Development and plasticity of the Drosophila larval neuromuscular junction. Wiley Interdiscip Rev Dev Biol 2:647–670.

Mosca TJ, Hong W, Dani VS, Favaloro V, Luo L (2012) Trans-synaptic Teneurin signalling in neuromuscular synapse organization and target choice. Nature 484:237–241.

Mosca TJ, Schwarz TL (2010) The nuclear import of Frizzled2-C by Importins-β11 and α2 promotes postsynaptic development. Nat Neurosci 13:935–943.

Nagarkar-Jaiswal S, Lee P-T, Campbell ME, Chen K, Anguiano-Zarate S, Gutierrez MC, Busby T, Lin W-W, He Y, Schulze KL, Booth BW, Evans-Holm M, Venken KJ, Levis RW, Spradling AC, Hoskins RA, Bellen HJ (2015) A library of MiMICs allows tagging of genes and reversible, spatial and temporal knockdown of proteins in Drosophila. Elife 4:e05338.

Neuert H, Deing P, Krukkert K, Naffin E, Steffes G, Risse B, Silies M, Klämbt C (2019) The Drosophila NCAM homolog Fas2 signals independently of adhesion. Development 147:dev181479.

Nievergelt CM et al. (2024) Genome-wide association analyses identify 95 risk loci and provide insights into the neurobiology of post-traumatic stress disorder. Nat Genet 56:792–808.

Noordermeer JN, Kopczynski CC, Fetter RD, Bland KS, Chen W-Y, Goodman CS (1998) Wrapper, a Novel Member of the Ig Superfamily, Is Expressed by Midline Glia and Is Required for Them to Ensheath Commissural Axons in Drosophila. Neuron 21:991–1001.

Nose A (2012) Generation of neuromuscular specificity in Drosophila: novel mechanisms revealed by new technologies. Front Mol Neurosci 5:62.

Özkan E, Carrillo RA, Eastman CL, Weiszmann R, Waghray D, Johnson KG, Zinn K, Celniker SE, Garcia KC (2013) An Extracellular Interactome of Immunoglobulin and LRR Proteins Reveals Receptor-Ligand Networks. Cell 154:228–239.

Öztürk-Çolak A et al. (2024) FlyBase: updates to the Drosophila genes and genomes database. GENETICS 227:iyad211.

Packard M, Koo ES, Gorczyca M, Sharpe J, Cumberledge S, Budnik V (2002) The Drosophila Wnt, Wingless, Provides an Essential Signal for Pre-and Postsynaptic Differentiation. Cell 111:319–330.

Parcerisas A, Ortega-Gascó A, Pujadas L, Soriano E (2021) The Hidden Side of NCAM Family: NCAM2, a Key Cytoskeleton Organization Molecule Regulating Multiple Neural Functions. Int J Mol Sci 22:10021.

Parcerisas A, Pujadas L, Ortega-Gascó A, Perelló-Amorós B, Viais R, Hino K, Figueiro-Silva J, Torre AL, Trullás R, Simó S, Lüders J, Soriano E (2020) NCAM2 Regulates Dendritic and Axonal Differentiation through the Cytoskeletal Proteins MAP2 and 14-3-3. Cereb Cortex 30:3781–3799.

Perry S, Han Y, Das A, Dickman D (2017) Homeostatic plasticity can be induced and expressed to restore synaptic strength at neuromuscular junctions undergoing ALS-related degeneration. Hum Mol Genet 26:4153–4167.

Perry S, Han Y, Qiu C, Chien C, Goel P, Nishimura S, Sajnani M, Schmid A, Sigrist SJ, Dickman D (2022) A glutamate receptor C-tail recruits CaMKII to suppress retrograde homeostatic signaling. Nat Commun 13:7656.

Petersen SA, Fetter RD, Noordermeer JN, Goodman CS, DiAntonio A (1997) Genetic Analysis of Glutamate Receptors in Drosophila Reveals a Retrograde Signal Regulating Presynaptic Transmitter Release. Neuron 19:1237–1248.

Petit F, Plessis G, Decamp M, Cuisset J-M, Blyth M, Pendlebury M, Andrieux J (2015) 21q21 deletion involving NCAM2: Report of 3 cases with neurodevelopmental disorders. Eur J Méd Genet 58:44–46.

Piechotta K, Dudanova I, Missler M (2006) The resilient synapse: insights from genetic interference of synaptic cell adhesion molecules. Cell Tissue Res 326:617–642.

Pielage J, Fetter RD, Davis GW (2005) Presynaptic Spectrin Is Essential for Synapse Stabilization. Curr Biol 15:918–928.

Polo-Parada L, Bose CM, Landmesser LT (2001) Alterations in Transmission, Vesicle Dynamics, and Transmitter Release Machinery at NCAM-Deficient Neuromuscular Junctions. Neuron 32:815–828.

Qin G, Schwarz T, Kittel RJ, Schmid A, Rasse TM, Kappei D, Ponimaskin E, Heckmann M, Sigrist SJ (2005) Four Different Subunits Are Essential for Expressing the Synaptic Glutamate Receptor at Neuromuscular Junctions of Drosophila. J Neurosci 25:3209–3218.

Qiu C, Perry S, Chen C, Chen J, Zhuang J, Han Y, Goel P, Dickman D (2025) Nonionic signaling rapidly remodels postsynaptic DLG to induce retrograde homeostatic plasticity. Proc Natl Acad Sci 122:e2502997122.

Rafuse VF, Polo-Parada L, Landmesser LT (2000) Structural and Functional Alterations of Neuromuscular Junctions in NCAM-Deficient Mice. J Neurosci 20:6529–6539.

Restrepo LJ, DePew AT, Moese ER, Tymanskyj SR, Parisi MJ, Aimino MA, Duhart JC, Fei H, Mosca TJ (2022) γ-secretase promotes Drosophila postsynaptic development through the cleavage of a Wnt receptor. Dev Cell.

Schmid A, Qin G, Wichmann C, Kittel RJ, Mertel S, Fouquet W, Schmidt M, Heckmann M, Sigrist SJ (2006) Non-NMDA-Type Glutamate Receptors Are Essential for Maturation But Not for Initial Assembly of Synapses at Drosophila Neuromuscular Junctions. J Neurosci 26:11267–11277.

Schuster CM, Davis GW, Fetter RD, Goodman CS (1996a) Genetic Dissection of Structural and Functional Components of Synaptic Plasticity. I. Fasciclin II Controls Synaptic Stabilization and Growth. Neuron 17:641–654.

Schuster CM, Davis GW, Fetter RD, Goodman CS (1996b) Genetic Dissection of Structural and Functional Components of Synaptic Plasticity. II. Fasciclin II Controls Presynaptic Structural Plasticity. Neuron 17:655–667.

Seeger M, Tear G, Ferres-Marco D, Goodman CS (1993) Mutations affecting growth cone guidance in drosophila: Genes necessary for guidance toward or away from the midline. Neuron 10:409–426.

Shen K, Scheiffele P (2010) Genetics and Cell Biology of Building Specific Synaptic Connectivity. Annu Rev Neurosci 33:473–507.

Shetty A, Sytnyk V, Leshchyns’ka I, Puchkov D, Haucke V, Schachner M (2013) The Neural Cell Adhesion Molecule Promotes Maturation of the Presynaptic Endocytotic Machinery by Switching Synaptic Vesicle Recycling from Adaptor Protein 3 (AP-3)-to AP-2-Dependent Mechanisms. J Neurosci 33:16828–16845.

Shiwaku H, Katayama S, Kondo K, Nakano Y, Tanaka H, Yoshioka Y, Fujita K, Tamaki H, Takebayashi H, Terasaki O, Nagase Y, Nagase T, Kubota T, Ishikawa K, Okazawa H, Takahashi H (2022) Autoantibodies against NCAM1 from patients with schizophrenia cause schizophrenia-related behavior and changes in synapses in mice. Cell Rep Med 3:100597.

Sink H, Rehm EJ, Richstone L, Bulls YM, Goodman CS (2001) sidestep Encodes a Target-Derived Attractant Essential for Motor Axon Guidance in Drosophila. Cell 105:57–67.

Sytnyk V, Leshchyns’ka I, Nikonenko AG, Schachner M (2006) NCAM promotes assembly and activity-dependent remodeling of the postsynaptic signaling complex. J Cell Biol 174:1071–1085.

Tang A-H, Chen H, Li TP, Metzbower SR, MacGillavry HD, Blanpied TA (2016) A trans-synaptic nanocolumn aligns neurotransmitter release to receptors. Nature 536:210–214.

Tessier-Lavigne M, Goodman CS (1996) The Molecular Biology of Axon Guidance. Science 274:1123–1133.

Wagh DA, Rasse TM, Asan E, Hofbauer A, Schwenkert I, Dürrbeck H, Buchner S, Dabauvalle M-C, Schmidt M, Qin G, Wichmann C, Kittel R, Sigrist SJ, Buchner E (2006) Bruchpilot, a Protein with Homology to ELKS/CAST, Is Required for Structural Integrity and Function of Synaptic Active Zones in Drosophila. Neuron 49:833–844.

Williams SE, Mealer RG, Scolnick EM, Smoller JW, Cummings RD (2020) Aberrant glycosylation in schizophrenia: a review of 25 years of post-mortem brain studies. Mol Psychiatry 25:3198–3207.

Yoshihara M, Rheuben MB, Kidokoro Y (1997) Transition from Growth Cone to Functional Motor Nerve Terminal inDrosophila Embryos. J Neurosci 17:8408–8426.

Yu M, Tan Q, Dong W, Xiang B (2025) Identification and prioritization of gene sets associated with schizophrenia risk by network analysis. Psychopharmacology:1–11.

Zhang B, Stewart B (2010) Electrophysiological Recording from Drosophila Larval Body-Wall Muscles. Cold Spring Harb Protoc 2010:pdb.prot5487.

Zito K, Parnas D, Fetter RD, Isacoff EY, Goodman CS (1999) Watching a Synapse Grow Noninvasive Confocal Imaging of Synaptic Growth in Drosophila. Neuron 22:719–729.

